# TRIM28 PRESERVES OVARIAN IDENTITY BY STABILIZING LINEAGE-SPECIFIC TRANSCRIPTION FACTOR HUBS

**DOI:** 10.64898/2026.07.21.739817

**Authors:** Laura Sitkiewicz, Florian Chaleil, Gaby Granès, Moïra Rossitto, Stéphanie Déjardin, Florence Cammas, Anne-Marie Lefrançois-Martinez, Isabelle Stévant, Francis Poulat

**Affiliations:** Institute of Human Genetics, CNRS UMR9002 University of Montpellier, 34396, Montpellier, France; Institut Génétique, Reproduction & Développement (iGReD), CNRS, INSERM, Université Clermont Auvergne, Clermont-Ferrand, France

## Abstract

Maintenance of ovarian cell identity is required throughout life to prevent the activation of the testicular program, but the epigenetic mechanisms underlying this process remain poorly understood. Although TRIM28 is required to prevent granulosa-to-Sertoli transdifferentiation, it can act both as a regulator of H3K9me3-dependent heterochromatin and as a transcriptional activator through its E3 SUMOligase activity. Here, we combined CUT&RUN, ATAC-seq and RNA-seq to define the respective contributions of these activities to maintain ovarian cell identity. Strikingly, only a small fraction of TRIM28-bound regions was associated with H3K9me3. Although *Trim28* deletion induced focal H3K9me3 loss, it had limited transcriptional consequences and primarily affected repetitive elements rather than regions controlling testis-determining genes. In contrast, *Trim28* loss led to reductions in chromatin accessibility and H3K27ac at regions enriched for ovarian transcription factor (TF) motifs FOXL2, NR5A2, ESR2 and RUNX1. Moreover, TRIM28 was frequently co-localized with these TFs on chromatin, and the accessibility and the SUMOylation at these co-bound regions were reduced by *Trim28* deletion. Together, our findings identify TRIM28 as a central organizer of ovarian TF hubs whose predominant function is to preserve granulosa cell identity through stabilization of lineage-specific TFs rather than H3K9me3-dependent heterochromatin.

**GRAPHICAL ABSTRACT:** 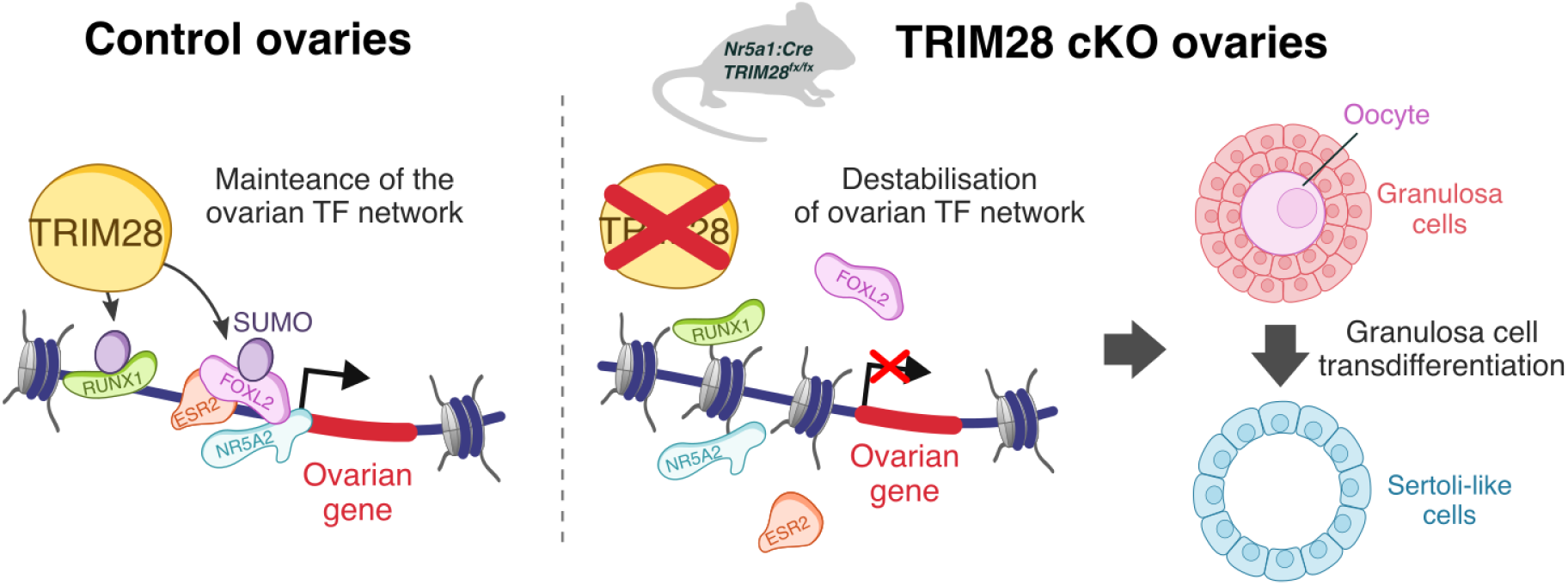

## INTRODUCTION

In vertebrates, gonadal sex is established during embryonic development. However, in some species, gonadal cell fate remains plastic throughout life. Gonads are composed of three principal cell lineages: germ cells, which give rise to the sex-specific gametes required for reproduction; supporting cells, which support germ cell development and differentiation (Sertoli cells in males and granulosa cells in females); and steroidogenic cells, which synthesize sex steroid hormones (Leydig cells in males and theca cells in females). Maintenance of supporting cell identity is required to preserve gonadal architecture, hormone production, germ cell support and fertility. When this maintenance is lost, ovarian tissue can partially or completely acquire male characteristics. Teleost fish provide a well-known example of this plasticity, as adult ovaries can naturally transform into testes through granulosa-to-Sertoli transdifferentiation in response to social or environmental cues (1). This shows that, in some vertebrates, the mechanisms maintaining granulosa and Sertoli cell identity remain reversible in adulthood.

In eutherian mammals, this plasticity is normally prevented by mechanisms that actively maintain ovarian identity after birth. The first evidence came from mice lacking both estrogen receptors (ESR1 and ESR2). Although ovarian differentiation proceeds normally during fetal development, adult ovaries progressively express the male transcription factor SOX9 and develop seminiferous tubule-like structures (2), with the severity of the phenotype depending on the genetic background (3). Later, conditional deletion of *Foxl2* showed that loss of this key ovarian transcription factor after birth activates *Sox9* expression and induces granulosa-to-Sertoli transdifferentiation, independently of oocyte loss (4). Conversely, ectopic expression of DMRT1 in granulosa cells is sufficient to induce a testicular fate (5, 6). Together, these studies demonstrate that ovarian identity requires continuous active maintenance throughout adult life.

We recently identified the chromatin regulator TRIM28 as an essential factor for the maintenance of ovarian identity (7). Conditional deletion of *Trim28* in fetal ovaries with the Nr5a1-Cre does not affect ovarian differentiation but induces progressive postnatal masculinization. Mutant ovaries lose FOXL2 expression in granulosa cells and progressively reorganize into seminiferous tubule-like structures expressing the Sertoli cell markers SOX8, SOX9 and DMRT1 (7). This phenotype closely resembles that observed after *Foxl2* deletion, suggesting that TRIM28 acts within the ovarian transcriptional network. However, how TRIM28 maintains ovarian identity remains unclear.

Although TRIM28 was initially identified as a transcriptional repressor (8–11), increasing evidence has shown that it can also activate gene expression (10, 12). Because TRIM28 does not bind DNA directly, it is recruited to chromatin by sequence-specific DNA-binding proteins, including KRAB zinc-finger proteins (KRAB-ZFPs) and other transcription factors (13, 14). In the ovary, TRIM28 cooperates with FOXL2 at ovarian regulatory elements to maintain the female transcriptional program while repressing testicular genes (7). Importantly, mice carrying a mutation in the TRIM28 PHD domain, which abolishes its SUMO E3 ligase activity, develop the same ovarian masculinization phenotype as *Trim28* conditional knockout mice, demonstrating that TRIM28-dependent SUMOylation (a post-translational modification affecting transcription factor activity) is required for maintenance of ovarian identity (7). Consistent with this, *Trim28*-deficient ovaries show major changes in chromatin SUMOylation, with loss of SUMOylation at ovarian regulatory regions and the appearance of de novo SUMOylation at regions associated with the male pathway (7).

TRIM28 regulates chromatin through at least two SUMO-dependent mechanisms. Through its SUMO E3 ligase activity, TRIM28 promotes SETDB1-dependent H3K9me3 deposition at transposable elements (TEs), contributing to their epigenetic silencing (9, 15) (Fig. 1a). TRIM28 also interacts with several transcription factors and regulates their activity through SUMOylation (16–18) (Fig. 1b). As such, the ovarian masculinization observed after *Trim28* deletion could result from defective H3K9me3-dependent heterochromatin maintenance, altered SUMOylation of ovarian transcription factors, or both. The relative contribution of these two SUMO-dependent functions of TRIM28 to the maintenance of ovarian identity remains unknown.

**Figure 1:**
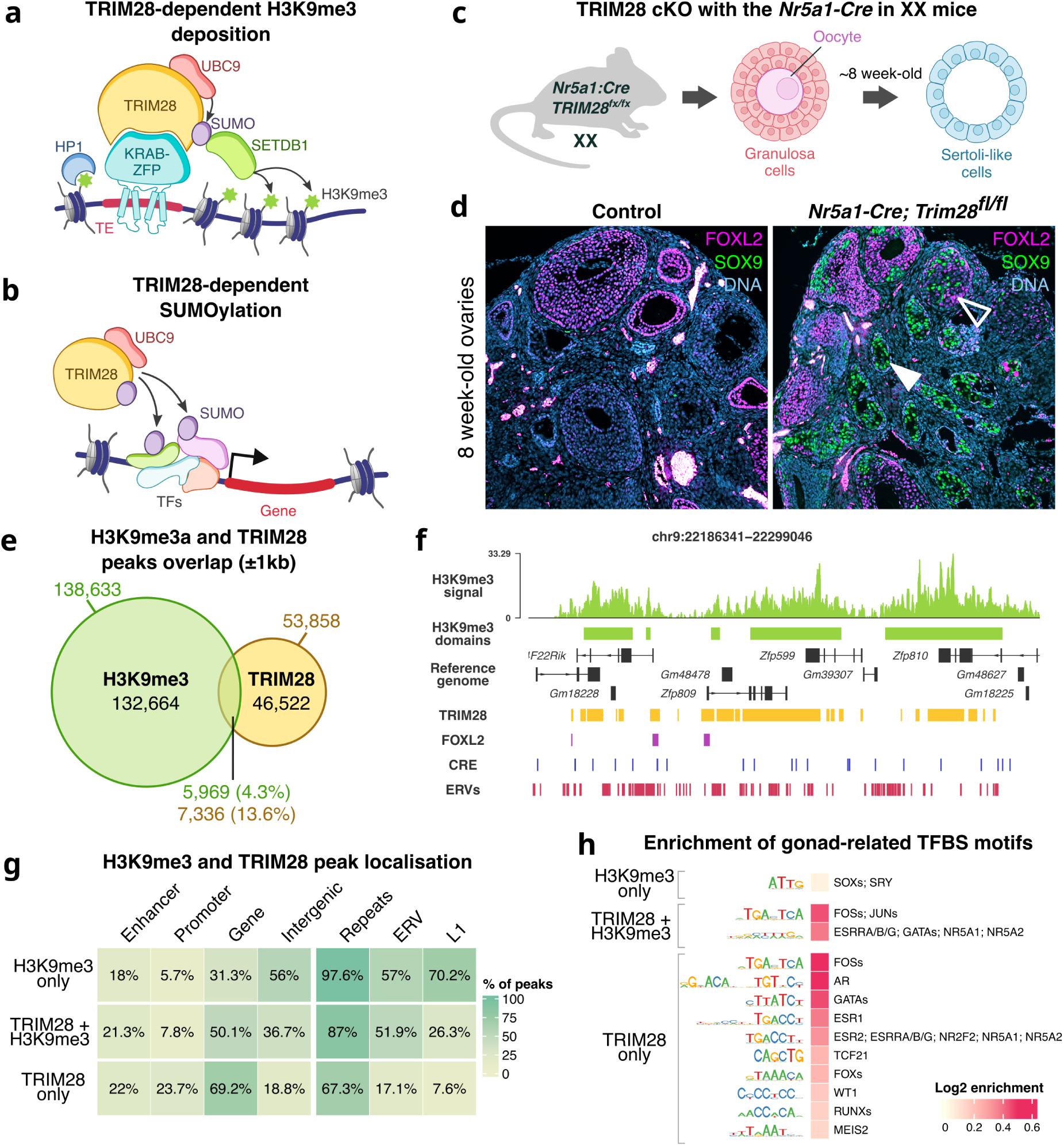
TRIM28 occupies distinct chromatin compartments in ovarian somatic cells. (**a**) Schematic of TRIM28’s canonical role in repressing transposable elements (TEs) via SETDB1-mediated H3K9me3 deposition and HP1 recruitment. (**b**) Schematic of TRIM28’s role in modulating transcription factor (TF) activity through SUMOylation, a post-translational modification that regulates TF stability, DNA binding, and cofactor recruitment. (**c**) Experimental model: Conditional deletion of *Trim28* in ovarian somatic cells using Nr5a1-Cre induces granulosa-to-Sertoli transdifferentiation. (**d**) Representative immunofluorescence of 8-week-old ovaries that shows FOXL2-positive granulosa cells (purple) in control ovaries, while *Trim28* cKO ovaries contain SOX9-positive Sertoli-like cells (green). Empty arrowhead: cortical follicle with FOXL2+SOX9+ cells (intermediate state); plain arrowhead: medullary follicle with SOX9+ only cells (transdifferentiated). (**e**) Venn diagram showing the overlap between TRIM28 binding sites and H3K9me3-enriched regions (±1 kb). (**f**) Genome browser track showing H3K9me3-enriched domains (green) in control ovaries, alongside TRIM28 and FOXL2 ChIP-seq peaks (orange and purple, respectively), ERVs (red bars), and cis-regulatory elements (CREs) (blue bars). (**g**) Heatmap showing the percentage of TRIM28-only, TRIM28+H3K9me3, and H3K9me3-only peaks overlapping genomic features (promoters, enhancers, gene bodies, repeats, ERVs, LINE1). The total sum is > 100% as a single peak can overlap several features simultaneously. (**h**) Transcription factor binding site (TFBS) motif enrichment in H3K9me3-only (top), TRIM28+H3K9me3 (middle) and TRIM28-only (bottom) peaks.

To determine which SUMO-dependent function of TRIM28 maintains ovarian identity, we combined ATAC-seq with H3K9me3 and H3K27ac CUT&RUN in control and *Trim28*-deficient ovaries, and integrated these datasets with TRIM28, FOXL2 and publicly available ChIP-seq profiles of ovarian transcription factors. Our results identify a previously unrecognized role for TRIM28 in organizing ovarian transcription factor regulatory hubs. Rather than acting primarily through H3K9me3-dependent heterochromatin maintenance, TRIM28 preserves ovarian identity by maintaining the activity of lineage-specific cis-regulatory elements.

## MATERIALS AND METHODS

### Mice and ethics

Animal care and handling were according to the “Réseau des Animaleries de Montpellier” (RAM) guidelines. Mouse lines were kept on a mixed 129SV/C57BL6/J background. Animals were housed on a 12/12 h light/dark cycle and maintained at a temperature of 22-24 °C, with humidity at 30–70%.

To generate ovarian somatic cells from 8-week-old control and *Trim28* cKO mice, the conditional *Trim28* allele was combined with the Nr5a1:Cre line, as previously described (7), and with the Ai14 tdTomato reporter allele (19) (MGI: J:155793). Heterozygous Trim28^flox/+^; Tg(Nr5a1:Cre) females displayed no detectable histological abnormalities and showed normal fertility and were used as control animals. For all experiments, ovaries from at least three females per genotype were collected, ensuring a minimum of three independent biological replicates.

### Genomic DNA isolation and transgenic mice genotyping

Genomic DNA was isolated from tail biopsies and PCR-amplified using the Platinum Direct PCR Universal Master Mix kit (Invitrogen, ref. A44647500).

The *Trim28* floxed allele was genotyped using a three-primer strategy as previously described (20): 5′-GGAATGGTTGTTCATTGGTG-3′, 5′-ACCTTGGCCCATTTATTGATAAAG-3′, and 5′-GCGAGCACGAATCAAGGTCAG-3′. This assay generates a 152 bp PCR product for the wild-type allele and a 180 bp product for the floxed allele. The Nr5a1:Cre transgene (21) was genotyped using the following primers: 5′-TCGGGGTTTTGTTCTCAGAC-3′ and 5′-ATGTTTAGCTGGCCCAAATG-3′, yielding a 500 bp PCR product. The Ai14-tdTomato reporter allele was genotyped using the primers 5′-CTGTTCCTGTACGGCATGG-3′ and 5′-GGCATTAAAGCAGCGTATCC-3′, producing a 196 bp amplicon specific for the mutant allele. The wild-type Rosa26 allele was detected using 5′-AAGGGAGCTGCAGTGGAGTA-3′ and 5′-CCGAAAATCTGTGGGAAGTC-3′ primers, generating a 297 bp PCR product. PCR conditions for all genotyping were: 94°C for 15 sec; 60°C for 15 sec, 68 °C for 20 sec, 35 cycles.

### Immunofluorescence staining

Ovaries were collected from 8-week-old mice and fixed overnight in 4% paraformaldehyde at 4°C. Immunofluorescence analyses were performed on 3.5 µm paraffin-embedded tissue sections. Sections were first dried on a heating plate, and paraffin was removed using Histo-Clear II (Electron Microscopy Sciences, 64111-01) with two successive 20-minute incubations. Samples were then rehydrated through a graded ethanol series (100% ethanol, 2 x 5 min; 75%, 50%, and 25% ethanol, 10 min each). Antigen retrieval was performed by incubating slides in 10 mM sodium citrate buffer (pH 6.9) for 20 minutes at 98°C using a Montage Opus365 apparatus (Diagnostic BioSystems). Sections were subsequently blocked for 1 hour at room temperature in PBS containing 10% donkey serum and 0.1% Triton X-100.

Primary antibodies (Supplementary Table S1) were applied overnight at 4°C in PBS supplemented with 0.1% Triton X-100 (see Table for dilutions). After washing, sections were incubated with Alexa Fluor–conjugated secondary antibodies for 30 minutes at room temperature. Slides were then mounted using Fluoroshield mounting medium with DAPI to visualize nuclei. Images were acquired using a Leica DM6B Thunder microscope.

### FACS isolation of Cre-activated Ai14 (tdTomato⁺) cells

Ovaries were collected from mice at 8-weeks old of each genotype and carrying Ai14 reporter. Ovaries subjected to CUT&RUN have been fixed with 0.1% paraformaldehyde in PBS for 1 minute and blocked with 12.5 % glycine for 5 minutes. Ovaries subjected to ATAC-seq were not fixed. Ovaries were washed three times with 1 mL of 1X PBS and digested in dissociation buffer: 1X HBSS + 10% FBS with 0.05mg/mL collagenase (Worthington Biochemical LS004176) for 1 hour at 37°C. Then digested ovarian cells were washed twice with 1X HBSS and isolated cells were selected on 30 µm pre-separation filters. TdTomato positive cells were purified using BD Melody FACS, centrifuged and frozen in FBS + 10% DMSO at -80°C.

### CUT&RUN and library prep

The CUT&RUN experimental protocol was adapted from the method described by Skene and Henikoff (22). The protocol was optimized for 100,000 cells per experiment and was performed in two independent experimental replicates for each genotype.

First, concanavalin A (ConA)-coated magnetic beads (BioMag®Plus Concanavalin A ref: 86057-3) were resuspended in bead activation buffer (20 mM HEPES-KOH, pH 7.9; 10 mM KCl; 1 mM CaCl₂; 1 mM MnCl₂). FACS-sorted ovarian somatic cells 0.1% PFA-fixed were thawed in a 37°C water bath for 5 minutes and collected to obtain 100,000 cells per tube. Cells were centrifuged at 800 g for 3 minutes and resuspended in wash buffer (20 mM HEPES-KOH, pH 7.9; 150 mM NaCl; 1% Triton X-100; 0.05% SDS; 0.5 mM spermidine supplemented with Roche Complete EDTA-free protease inhibitor) for two washes. Then, 20 µL of activated beads were added to each tube containing the cells in wash buffer. To preserve cell integrity, tubes were placed on a magnetic rack to remove the wash buffer, and antibodies (Supplementary Table S1) were added at a 1:100 dilution in Antibody Buffer (20 mM HEPES-KOH, pH 7.9; 150 mM NaCl; 1% Triton X-100; 0.05% SDS; 2 mM EDTA; 0.05% digitonin). Cells were incubated with rotation overnight at 4°C.

After removal of the antibody buffer, cells were washed twice with permeabilization buffer (20 mM HEPES-KOH, pH 7.9; 150 mM NaCl; 1% Triton X-100; 0.05% SDS; 0.5 mM spermidine; 0.05% digitonin supplemented with Roche Complete EDTA-free protease inhibitor). Cells were then resuspended in permeabilization buffer and incubated for 10 minutes at room temperature with 2.5 µL of pAG-MNase (20X-15-1016, EpiCypher). Cells were washed twice again with permeabilization buffer and resuspended in 100 µL of permeabilization buffer supplemented with 2 µL of 100 mM CaCl₂, followed by incubation for 2 hours at 4°C to allow MNase-mediated cleavage of target chromatin regions. Reactions were stopped by adding 100 µL of 2x stop buffer (340 mM NaCl; 20 mM EDTA; 4 mM EGTA; 0.02% digitonin; 0.25 mg/mL glycogen; 0.06 mg/mL RNase A) and incubating at 37°C for 15 minutes.

Cells and beads were then captured using a magnetic rack, and the supernatant containing CUT&RUN DNA fragments was collected. De-crosslinking was performed by incubating samples at 55°C for 4 hours in the presence of 2 µL of 10% SDS and 3 µL of 20 mg/mL proteinase K. CUT&RUN DNA fragments were purified by phenol–chloroform extraction. DNA precipitation was carried out at -80°C in ethanol in the presence of 3.5 M sodium acetate (pH 5.2) and glycogen. DNA was then resuspended in 20 µL of Tris-HCl (pH 8.5) to generate CUT&RUN libraries using the NEBNext Ultra II DNA Library Prep Kit for Illumina and NEBNext Multiplex Oligos for Illumina. Libraries were sequenced as 150 bp paired-end reads on an Illumina NovaSeq X Plus system (Novogene).

### ATAC-seq and library prep

To perform ATAC-seq experiments, we used the Active Motif ATAC-Seq Kit (Ref 53150). Experiments were carried out in three independent biological replicates for each genotype. Unfixed FACS-sorted ovarian somatic cells were first placed in a 37°C water bath for 5 minutes and then collected to obtain 75,000 cells per sample. Cells were centrifuged at 800 g and washed twice with cold 1x PBS. After the second wash, cells were resuspended in lysis buffer and centrifuged to isolate nuclei. Tagmentation was then performed by resuspending the nuclei in Tagmentation Master Mix and incubating the samples in a thermomixer for 30 minutes at 37°C with shaking at 800 rpm. Following tagmentation, 3 M sodium acetate and DNA Purification Binding Buffer were added, and DNA was purified using a column-based method. The tagmented DNA was subsequently amplified by PCR using indexed primers to generate libraries suitable for Illumina sequencing. PCR conditions for ATAC-seq library preparation were as follows: 72°C for 5 minutes; 98°C for 30 seconds; 63°C for 30 seconds; and 72°C for 1 minute, for a total of 10 cycles. ATAC-seq libraries were sequenced as 150 bp paired-end reads on an Illumina NovaSeq X Plus system (Novogene).

### CUT&RUN mapping, peak calling and differential enrichment analysis

H3K9me3 and H3K27ac CUT&RUN FASTQ files were processed using the nf-core/cutandrun pipeline (v3.2.2). Read quality was assessed with FastQC, adaptor sequences were removed using Cutadapt, and reads were aligned to the mouse mm10/GRCm38 genome assembly (Gencode M25) using Bowtie2. Reads mapping to ENCODE blacklist regions were excluded using the default nf-core parameters.

Peaks were called with SEACR using stringent settings (--seacr_stringent stringent), fragment extension (--extend_fragments TRUE) and matched controls (--use_control TRUE). For each histone mark, a consensus peak set was generated by merging peaks detected in at least one replicate. H3K9me3 domains were defined by merging adjacent peaks separated by < 2 kb using bedtoolsr (v2.30.0-7).

Fragment counts per peak were quantified using featureCounts (v2.1.1) from paired-end aligned reads. All downstream analyses were performed in R (v4.6.0). Peaks with < 50 fragments across all samples were discarded. Differential enrichment analysis was performed with DESeq2 (v1.52.0) using raw fragment counts. Peaks with adjusted P-values < 0.01 and | log2FC| > 0.5 were considered significantly differentially enriched.

Variance stabilizing transformation (VST) from DESeq2 was used for data visualization and clustering analyses. BigWig files were normalized using DESeq2 size factors, and replicate tracks from the same condition were merged for visualization. Genomic tracks were generated using Gviz (v1.56.0), and coverage heatmaps were produced with EnrichedHeatmap (v1.42.0).

### ATAC-seq mapping, peak calling and differential accessibility analysis

ATAC-seq FASTQ files were processed using the nf-core/atacseq pipeline (v2.1.2). Read quality was assessed with FastQC, adaptor sequences were removed using Cutadapt, and reads were aligned to the mouse mm10/GRCm38 genome assembly (Gencode M25) using BWA. Mapped reads were filtered using the default nf-core parameters.

Accessible chromatin regions were identified with MACS2 using narrow peak calling (--narrow_peak TRUE) and a false discovery rate threshold of 0.01 (--macs_fdr 0.01). A consensus peak set was generated by retaining peaks detected in at least two replicates. Fragment counts per peak were quantified using featureCounts from paired-end aligned reads. Peaks with fewer than 20 fragments across all samples were discarded.

Differential chromatin accessibility analysis was performed in R (v4.6.0) using DESeq2 (v1.52.0) on raw fragment counts. Peaks with adjusted P-values < 0.01 and |log2FC| > 0.5 were considered significantly differentially accessible. BigWig files were normalized using DESeq2 size factors, and replicate tracks from the same condition were merged for visualization. Heatmaps of ATAC-seq signal centered on peak regions were generated from the merged normalized BigWig files using EnrichedHeatmap.

### ChIP-seq mapping and peak calling

Publicly available TRIM28 (7), FOXL2 (7) (GEO ref. GSE166383) NR5A2 (a.k.a. LRH-1, control ovaries only) (23, 24) (GEO ref. GSE119508 and GSE208106), ESR2 (6, 25) (GEO ref. GSE203391 and GSE154484), and RUNX1 (control ovaries only, GEO ref. GSE152941) (26) ChIP-seq datasets were reanalyzed uniformly. FASTQ files were retrieved from the Gene Expression Omnibus (GEO) database using nf-core/fetchngs (v1.12.0) and processed with the nf-core/chipseq pipeline (v2.0.0). Read quality was assessed with FastQC, adaptor sequences were removed using Cutadapt, and reads were aligned to the mouse mm10/GRCm38 genome assembly (Gencode M25) using BWA. Mapped reads were filtered using the default nf-core parameters.

Peaks were called with MACS2. Both broad and narrow peak calling modes were initially tested, and the resulting peaks and coverage tracks were visually inspected to determine the most appropriate setting for each dataset. TRIM28, FOXL2, NR5A2 and ESR2 datasets were analyzed using broad peak calling, as narrow peak calling failed to identify visually supported binding regions. In contrast, RUNX1 peaks were called using narrow peak mode because broad peak calling generated extensive non-specific enrichment. NR5A2 and ESR2 peaks from the different studies were merged to obtain a larger peak dataset.

### Peak annotations and overlaps

CUT&RUN, ATAC and ChIP-seq peak annotation on general genomic features (promoters, intron, exon, intergenic, nearest gene) was done using ChIPseeker (v1.48.0) with the GTF file used for mapping (from Gencode, M25). Peak annotation over CRE/enhancers and repeats was done by merging mm10 enhancer annotation from FANTOM5 and ENCODE cCRE databases, and by using the mm10 RepeatMasker database, respectively.

Peak overlaps between two datasets were computed with GenomicRanges and visualized as pie chat with ggplot2. Overlaps with more than two datasets represented as Venn diagrams and Upset plot were computed with gVenn (v1.1.1). The numbers with the peak overlap tables and the gVenn representations may differ. This is due to the difference in the overlap calculation method. In gVenn, overlaps were computed on non-redundant genomic regions obtained by merging all intervals across datasets.

### TFBS motif analysis

Transcription factor binding site (TFBS) motif enrichment analysis was performed using monaLisa (v1.18.0) with vertebrate TFBS position weight matrices from the JASPAR2024 database retrieved using TFBSTools (v1.50.0). Full peak regions were used for motif scanning. Enrichment analyses were performed against a random genomic background matched for GC content. A stringent significance threshold of FDR < 10^-10^ was applied to minimize false positive enrichments.

To facilitate interpretation and visualization, significantly enriched motifs were clustered according to motif similarity using motifStack with cutoffPval = 0.001. Results were represented as heatmaps showing the mean log2 enrichment of grouped motifs together with consensus sequence logos and associated transcription factor names using ComplexHeatmap.

### GO term enrichment analyses

GO term enrichment analyses were done using clusterProfiler (v4.20.0).

### Single-cell RNA-seq expression profiles

Single-cell RNA-seq data were obtained from (7). Data were loaded in Seurat (v5.1.0) using the gene, barcode, expression matrix and metadata provided on GEO. Raw count matrices from three control and three mutant ovaries were imported and subjected to standard quality filtering (≥500 detected genes per cell and <20% mitochondrial transcripts). Cell identities were assigned based on the published annotations and marker gene expression. For differential expression analyses, raw UMI counts were aggregated by biological replicate using AggregateExpression to generate pseudobulk samples for each cell population. Differential expression was performed with DESeq2 (design ∼ group) after filtering genes with fewer than 10 counts in at least two samples. Comparisons were performed between *Trim28* cKO and control ovaries.

### Proximity ligation assay

Ovaries were collected and fixed in 4% PFA overnight at 4 °C and were then washed in PBS and stored at 4 °C in 70% ethanol and posteriorly paraffin embedded. 5 μm paraffin sections were dewaxed, rehydrated and subjected to antigen retrieval in boiling citrate sodium buffer (10mM, 0,05% tween20) (P1379, Sigma-Aldrich) for 20mn. Sections were blocked for an hour with PBS containing 2.5% horse serum (Vector Laboratories) and incubated overnight at 4 °C with primary antibody (Supplementary Table S1). After washing, slides were incubated with ImPRESS secondary antibodies (ImPress Polymer Detection Kit, Vector Laboratories) for 1h at room temperature. Polymer-coupled horseradish peroxidase (HRP) activity was then detected with tyramide signal amplification (TSA)-Alexa–coupled fluorochromes for fluorescence (Thermo Fisher Scientific, Alexa_488 B40953, Alexa_555 B40955). Slides were then mounted with Vectashield Hardset solution containing DAPI (Vector Laboratories). Antibodies are listed in Table n. At least three ovaries of each genotype were analyzed for each marker.

For proximity ligation assay (PLA), previously blocked sections were incubated overnight at 4 °C with indicated antibodies followed by Duolink in situ PLA (Sigma-Aldrich) anti-rabbit (plus) probes and anti-mouse (minus) or anti-goat (minus) detection reagents according to manufacturer’s instructions. Images were acquired with a Zeiss AxioImager with Apotome2 using ZEN 3.11 blue edition software. Image settings and processing were identical across genotypes.

PLA signals were quantified with QuPath 0.6.0 software. Statistics were conducted using R. Normality of population distribution was assessed with Kolmogorov–Smirnov normality test. To compare two populations, unpaired, two-tailed t test was used for normally distributed data with the same variance or Mann–Whitney for non-normal distributions.

## RESULTS

### TRIM28 occupies repressive and active chromatin compartments to maintain ovarian cell identity

To investigate the relationship between TRIM28 binding and H3K9me3 in ovarian somatic cells, we first profiled the genomic distribution of H3K9me3, a hallmark of heterochromatin that is deposited by SETDB1 in the canonical TRIM28 pathway (Fig. 1a). To specifically analyze the somatic cell population targeted by Nr5a1-Cre, tdTomato-positive cells from Nr5a1-Cre;tdTomato ovaries were purified by FACS prior to genomic analyses, thereby excluding germ cells. To ensure consistency with our previous study (7), all analyses were performed on 8-week-old ovaries, when granulosa-to-Sertoli transdifferentiation is ongoing but not yet complete. The medulla is largely reorganized into SOX9-positive pseudo-tubules, whereas cortical follicles still contain FOXL2-positive granulosa cells together with cells initiating SOX9 expression (Fig. 1c-d). As Cre activity is initiated during fetal stages and recruitment of the second follicular wave is completed shortly after birth (27, 28), chromatin changes associated with TRIM28 loss should already be largely established at this stage.

CUT&RUN analyses identified 138,633 H3K9me3 peaks, which merged into 87,759 enriched domains (peaks separated by <2 kb), with an average length of 5.4 kb (Supplementary Fig. S1, Supplementary Table S2) and covering approximately 18% of the mouse genome. Most H3K9me3 domains were intergenic (61.7%), while 24.8% overlapped introns and 7.9% were located on gene promoters (within 2 kb upstream of the transcription start sites; Supplementary Fig. S1d).

To assess the extent to which TRIM28 is associated with H3K9me3 in ovarian somatic cells, we compared these regions with TRIM28 and FOXL2 binding sites by reanalyzing ChIP-seq data from our previous study in 8-week-old control ovaries (7) (Supplementary Table S3). Given that H3K9me3 can spread from TRIM28 initial recruitment site, overlaps were assessed within a ±1kb window surrounding H3K9me3 peaks (15, 29). We found that 13.6% (7,336) of TRIM28 peaks overlapped or were located within ±1 kb of a H3K9me3-enriched region (Fig. 1e, Supplementary Table S2) and one third of them (1,794) overlapping also with FOXL2 (Supplementary Fig. S1e). As an example, we observed that TRIM28 co-localized with H3K9me3 at ZFP genes (Fig. 1f), consistent with previous reports (30, 31). Conversely, only 4.3% of H3K9me3 peaks overlapped TRIM28 binding sites. These results indicate that, in the adult ovary, TRIM28 is directly associated with only a limited fraction of the H3K9me3 chromatin landscape, suggesting that it regulates a restricted subset of H3K9me3-enriched regions.

We next compared the genomic features associated with H3K9me3-only, TRIM28-associated H3K9me3 and TRIM28-only regions using a non-exclusive overlap analysis (i.e. each region could overlap multiple genomic features simultaneously, and each of them is counted) (Fig. 1g). Almost all H3K9me3-only regions overlapped with repetitive sequences (97.6%), particularly L1 elements (70.2%), consistent with the repeat-rich nature of heterochromatin. TRIM28-associated H3K9me3 regions also overlapped extensively repetitive elements (87%) but were more frequently associated with gene bodies (50.1%) than H3K9me3-only regions (31.3%). In contrast, TRIM28-only peaks were more found at promoters (23.7%) and gene bodies (69.2%) and showed a lower overlap with repetitive elements (67.3%). These analyses indicate that TRIM28 occupies two distinct chromatin compartments in ovarian somatic cells: canonical H3K9me3-rich heterochromatin associated with repetitive sequences, and gene regulatory regions that are largely independent of H3K9me3.

Because TRIM28 does not bind DNA directly but is recruited to chromatin by sequence-specific DNA-binding proteins, we compared transcription factor binding site (TFBS) motif enrichment between TRIM28-only peaks and TRIM28+H3K9me3 peaks. We identified 361 significantly enriched TFBS motifs in TRIM28-only peaks, including motifs recognized by key gonadal and ovarian transcription factors such as GATA4, ESR1/2, FOXL2, FOS, NR5A1, NR5A2, RUNX1 and WT1 (Fig. 1h, Supplementary Table S4). In contrast, only 35 motifs were enriched in TRIM28+H3K9me3 regions. These included FOS, GATA4, NR5A1 and NR5A2, suggesting that recruitment of TRIM28 to both euchromatin and heterochromatin can involve gonadal transcription factors. H3K9me3-only regions showed a different motif landscape, with enrichment for ZFP-related motifs (Supplementary Table S4), consistent with their transposable element-rich composition (32). Apart from a modest enrichment for SOX/SRY motifs (Fig. 1h, Supplementary Table S4), no enrichment for gonadal TFBS was detected in the H3K9me3-only regions.

Together, these results reveal two distinct chromatin compartments associated with TRIM28 in ovarian somatic cells: a canonical H3K9me3-rich heterochromatin compartment and a larger regulatory compartment enriched for ovarian transcription factor binding sites. These findings indicate that TRIM28 is associated not only with heterochromatin organization but also with potential ovarian cis-regulatory elements, suggesting two mechanistically distinct functions in the maintenance of ovarian identity.

### Focal loss of H3K9me3 following TRIM28 loss does not target testis-determining loci

Given that loss of TRIM28 in ovarian somatic cells induces granulosa-to-Sertoli-like transdifferentiation, and that TRIM28 promotes heterochromatin formation through H3K9me3 deposition, we asked whether testis-determining genes or their regulatory elements are embedded within TRIM28-associated heterochromatin and become aberrantly activated upon its loss. To address this, we profiled genome-wide changes in H3K9me3 distribution following *Trim28* deletion using the Nr5a1-Cre (hereafter referred to as *Trim28* cKO). H3K9me3 profiles were generated as before by CUT&RUN from purified somatic cells of 8-week-old cKO ovaries and compared to H3K9me3 profiles from control somatic cells (Supplementary Fig. S2, Supplementary Table S2).

In the absence of TRIM28, 4,127 genomic regions showed a significant reduction in H3K9me3 signal compared with controls, representing only 3% (4,127 out of 138,633) of all H3K9me3 regions detected in control ovaries (Fig. 2a, Supplementary Table S5), whereas 1,708 regions showed increased H3K9me3 (Supplementary Fig. S3, Supplementary Table S5). 35.3% of H3K9me3-reduced regions in *Trim28* cKO ovaries were located within ±1 kb of a TRIM28 binding site identified in controls (Fig. 2b). This proportion was greater than expected from the genome-wide distribution of TRIM28-associated H3K9me3 regions (Fisher’s exact test, odds ratio = 15.7, p < 2.2×10^-^¹⁶), indicating that H3K9me3 loss occurs preferentially at TRIM28-associated heterochromatin. On the other hand, the vast majority (74.7%) of all TRIM28-associated H3K9me3 regions found in controls remained unchanged following *Trim28* deletion, indicating that TRIM28 is required for maintenance of only a subset of these domains.

**Figure 2:**
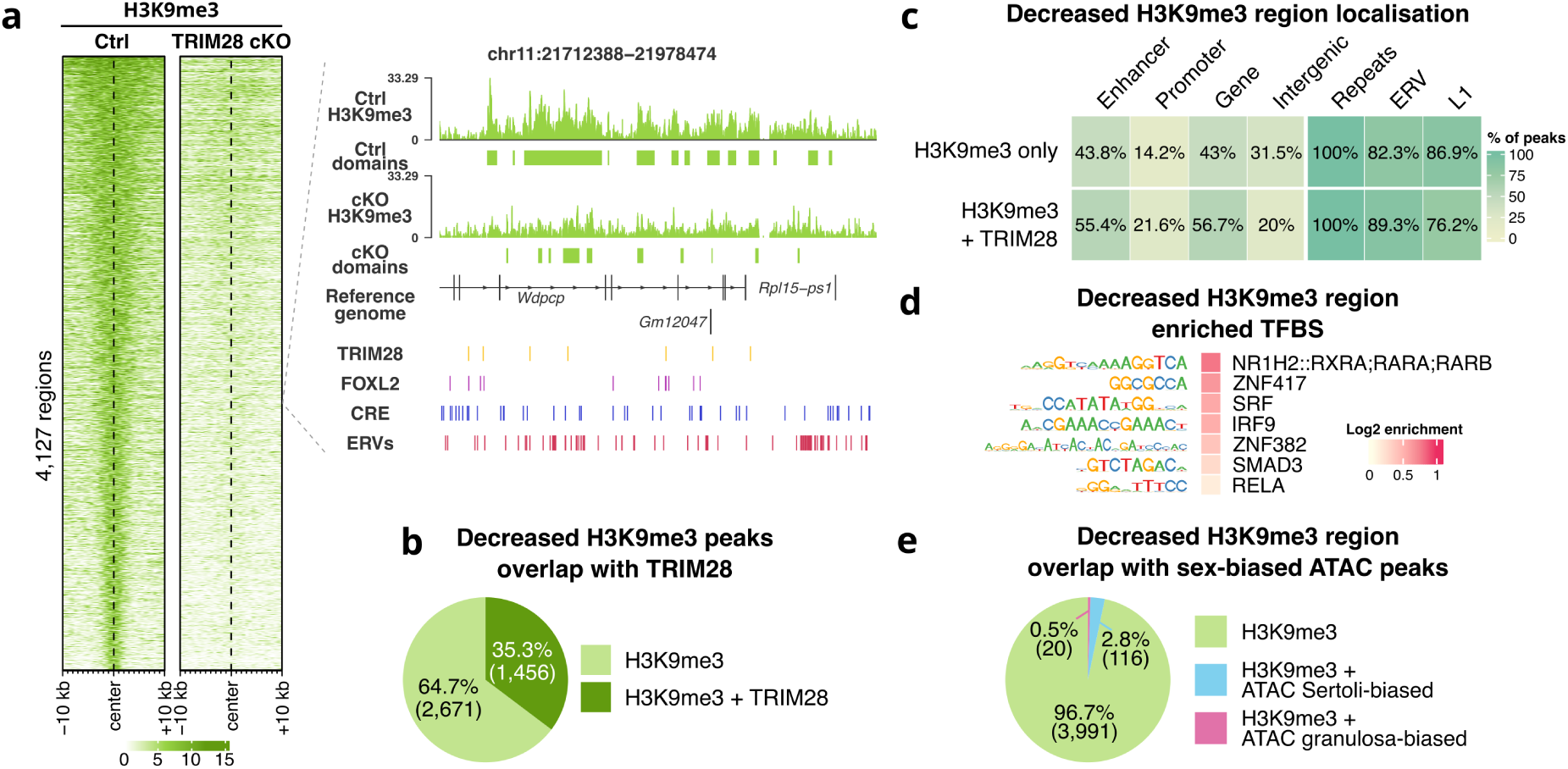
TRIM28 loss induces focal H3K9me3 remodeling without derepressing testicular genes. (**a**) Heatmap of H3K9me3 signal at 4,127 regions losing H3K9me3 in Trim28 cKO ovaries (vs. control). The dashed line indicates the center of the peaks. Right panel: Genome browser track of a representative region showing H3K9me3 loss in cKO (vs. control) and overlap with TRIM28 binding, FOXL2 ChIP-seq, CREs, and ERVs. (**b**) Pie chart showing the proportion of H3K9me3-decreased regions overlapping TRIM28 binding sites (±1 kb) in control ovaries. (**c**) Heatmap showing the percentage of H3K9me3-decreased regions overlapping genomic features (enhancers, promoters, gene bodies, intergenic regions, repeats, ERVs, LINE1). (**d**) TFBS motif enrichment in H3K9me3-decreased regions. All enriched motifs are shown. (**e**) Pie chart showing the overlap between H3K9me3-decreased regions and sex-biased open chromatin regions (from Lindeman et al., 2021). Sertoli-biased regions are colored in blue and granulosa-biased regions in pink.

As TRIM28-dependent H3K9me3 has been shown to spread over several tens of kilobases from TRIM28-bound loci *in vitro* (29), we asked whether H3K9me3-loss regions lacking direct TRIM28 binding could be explained by this mechanism. Of the 2,671 H3K9me3-loss regions not directly bound by TRIM28 in control ovaries, 593 (22%) were located within 50 kb of a TRIM28-associated H3K9me3-loss site, consistent with the reported spreading mechanism. The remaining regions were located >50 kb away, often several megabases from the nearest TRIM28-associated site, suggesting that their H3K9me3 loss may reflect indirect consequences of *Trim28* deletion. Conversely, regions gaining H3K9me3 rarely overlapped TRIM28 binding sites (3.3%), also suggesting indirect chromatin remodeling (Supplementary Fig. S3b).

Genomic annotation revealed that TRIM28-associated H3K9me3 loss affected slightly more gene regulatory regions, including enhancers (55.4%) and promoters (21.6%), compared to TRIM28-independent H3K9me3 loss (43.8M and 14.2%, respectively) (Fig. 2c). We identified a total of 1,418 genes in the regions with reduced H3K9me3. These genes showed only one enriched GO term (“homophilic cell-cell adhesion”) and did not include known testis-determining factors, except for the *Nr5a1* gene promoter (Supplementary Table S5). However, because the *Trim28* cKO model carries the *Nr5a1-Cre* transgene (with potentially several copies), we don’t know whether this signal originates from the endogenous locus or the transgene. Moreover, *Nr5a1* expression remained unchanged despite the local loss of H3K9me3, both in our single-cell RNA-seq analysis of 8-week-old cKO ovaries and in our bulk RNA-seq analysis of 7-month-old cKO ovaries (7). We therefore did not interpret this region further.

Among the genes overlapping with reduced H3K9me3, only 29 (2%) of them were transcriptionally upregulated in 8-week-old cKO ovaries (single-cell RNA-seq data, Supplementary Table S5) (7), and 134 (9.4%) in 7-month-old cKO ovaries (bulk RNA-seq data, Supplementary Table S5) (7), indicating that the decrease of H3K9me3 is not always sufficient to drive transcription activation.

To determine whether regions losing H3K9me3 contain testicular regulatory elements, we first performed transcription factor binding site motif enrichment analysis. Only nine motifs were significantly enriched in H3K9me3-loss regions (Fig. 2d, Supplementary Table S5), none of which corresponded to known testicular transcription factors. We next compared these regions with publicly available datasets of Sertoli- and granulosa-biased open chromatin regions (6). We found that only 116 regions (3%) overlapped Sertoli-biased regulatory elements (Fig. 2e, Supplementary Table S5), and none were located near known testis-determining genes. Together, these results indicate that regions losing H3K9me3 are unlikely to represent cryptic Sertoli regulatory elements responsible for granulosa-to-Sertoli-like transdifferentiation.

Together, these results show that TRIM28 loss induces focal depletion of H3K9me3, predominantly at repeat-rich genomic regions. However, these regions are not associated with activation of testis-determining genes, enrichment for Sertoli transcription factor motifs, or Sertoli-biased regulatory elements. These findings indicate that H3K9me3-dependent heterochromatin remodeling is unlikely to represent the main mechanism underlying granulosa-to-Sertoli-like transdifferentiation following *Trim28* deletion.

### Cis-regulatory network reprogramming reflects a switch from ovarian to Sertoli identity

Because H3K9me3 remodeling alone does not explain activation of the male transcriptional program following *Trim28* deletion, we next investigated whether TRIM28 loss affects chromatin accessibility and the activity of cis-regulatory regions. To this end, we performed ATAC-seq (Supplementary Fig. S4) and H3K27ac CUT&RUN (Supplementary Fig. S5) on purified somatic cells from control and *Trim28* cKO 8-week-old ovaries.

Genome-wide analysis identified significant changes in chromatin accessibility at 18.2% of ovarian somatic cell ATAC peaks (8,609 of 47,341, Fig. 3a, Supplementary Table S6). Among these differentially accessible regions (DARs), 3,187 showed reduced accessibility (labeled as “a”), whereas 5,422 gained accessibility in the *Trim28* cKO (labeled as “b”). Around 60% of regions losing accessibility (cluster a) overlapped TRIM28 binding sites in control ovaries (Fig. 3b), suggesting a direct role for TRIM28 in maintaining open chromatin, while 46.5% of regions gaining accessibility (cluster b) were also TRIM28-bound, indicating that TRIM28 can additionally restrain accessibility at specific loci (Fig. 3b).

**Figure 3:**
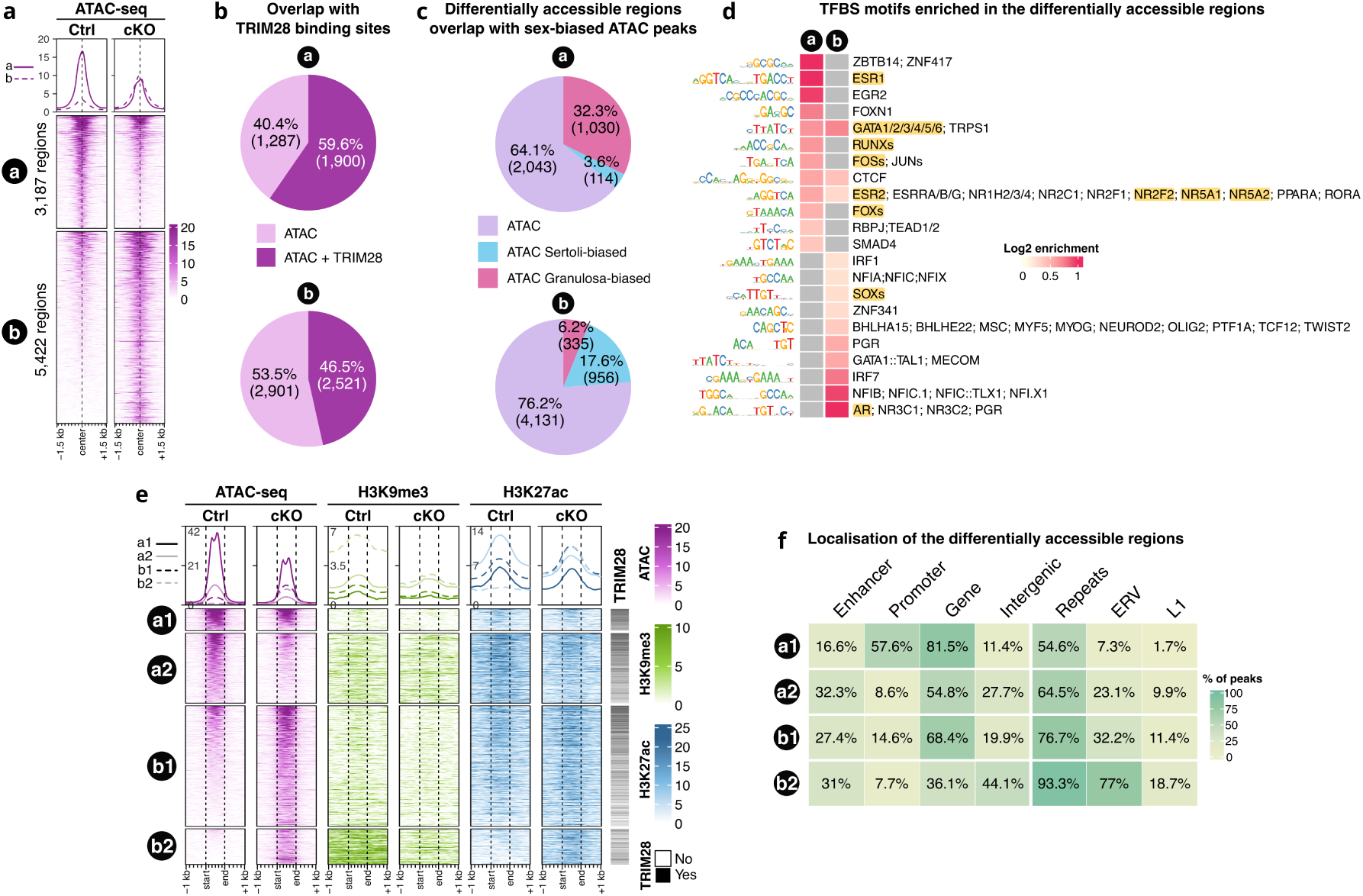
TRIM28 loss induces granulosa-to-Sertoli chromatin reprogramming. (**a**) Heatmap of ATAC-seq signal at 8,609 differentially accessible regions (DARs) in *Trim28* cKO vs. control ovaries. Cluster a (3,187 regions): loss of accessibility; Cluster b (5,422 regions): gain of accessibility. (**b**) Pie charts showing the overlap between DARs and TRIM28 binding sites in control ovaries. (**c**) Pie charts showing the proportion of DARs overlapping sex-biased open chromatin regions (from Lindeman et al., 2021). (**d**) TFBS motif enrichment in DARs. Grey squares: nonsignificant enrichments. (**e**) Hierarchical clustering of DARs based on H3K9me3 and H3K27ac coverage in control and *Trim28* cKO ovaries. Right bar: Presence of TRIM28 binding sites in each cluster (black = yes, white = no). (**f**) Heatmap showing the percentage of DARs overlapping genomic features (enhancers, promoters, gene bodies, repeats, ERVs, LINE1) for each cluster.

To determine whether changes in chromatin accessibility reflect the granulosa-to-Sertoli-like transdifferentiation observed in *Trim28* cKO ovaries, we compared DARs with previously identified granulosa- and Sertoli-biased open chromatin regions (6). We found that 1,030 (32.3%) regions losing accessibility corresponded to granulosa-biased regions (Fig. 3c, Supplementary Table S6), whereas 956 (17.6%) regions gaining accessibility corresponded to Sertoli-biased regions. These results indicate a progressive loss of the granulosa cis-regulatory program together with the establishment of a Sertoli-like regulatory landscape following TRIM28 loss.

Consistent with these observations, TFBS motif enrichment analysis revealed distinct transcription factor signatures in the two classes of DARs (Fig. 3d, Supplementary Table S6). Regions losing accessibility (cluster a) were enriched for motifs recognized by granulosa cell transcription factors, including NR5A2, FOXL2, ESR1/2 and RUNX1. In contrast, regions gaining accessibility (cluster b) were enriched for SOX family and androgen receptor (AR) motifs, consistent with activation of a Sertoli-like regulatory program. Unlike the granulosa-to-Sertoli transdifferentiation induced by ectopic DMRT1 expression (5), however, we did not detect enrichment of DMRT1 motifs in regions gaining accessibility. This is consistent with the low expression of *Dmrt1* in 8-week-old *Trim28* cKO ovaries (7).

To determine how changes in chromatin accessibility relate to histone mark remodeling following *Trim28* deletion, we integrated the DARs with H3K9me3 and H3K27ac CUT&RUN profiles from control and *Trim28* cKO ovaries. Hierarchical clustering identified four groups based on coordinated changes in chromatin accessibility and histone modifications (Fig. 3e, Supplementary Table S6). Among regions losing accessibility, cluster a1 (757 regions) showed decreased accessibility with only minor changes in H3K9me3 and H3K27ac. In contrast, cluster a2 (2,430 regions) showed a marked reduction in H3K27ac, while H3K9me3 remained largely unchanged. Thus, decreased chromatin accessibility following *Trim28* deletion is primarily associated with loss of H3K27ac rather than gain of the repressive H3K9me3 mark.

Regions gaining accessibility following TRIM28 loss were also subdivided into two clusters, b1 and b2. The largest cluster (b1; 4,185 regions) showed little change in H3K9me3 but acquired the active chromatin mark H3K27ac. In contrast, cluster b2 (1,237 regions) underwent the most extensive chromatin remodeling, with a marked loss of H3K9me3 together with increased chromatin accessibility and acquisition of H3K27ac following *Trim28* deletion (Fig. 1a).

Overlap with the repeatMasker annotation revealed that the b2 cluster was particularly rich in repeated sequences (93.3%, Fig. 3f), with 77% of the peaks overlapping at least one ERV. We examined whether the TE-derived sequences within these regions became transcribed following *Trim28* cKO. Despite their high abundance, only 189 of the 1,506 TE-derived sequences were overexpressed (33), the vast majority of which belonged to the ERV family (181; Supplementary Table S6). Because activated transposable elements can serve as alternative promoters or enhancers for nearby genes (34, 35), we examined whether the reactivated ERVs were located in the vicinity of key testis-determining genes (Supplementary Table S6). However, we found no evidence supporting such a mechanism in *Trim28 cKO* ovaries.

Together, these results show that TRIM28 loss induces widespread changes in chromatin accessibility and H3K27ac that are accompanied by changes in transcription factor motif composition, reflecting the progressive replacement of the ovarian cis-regulatory program by a Sertoli-like regulatory landscape. In contrast, newly accessible chromatin regions accompanied by H3K9me3 loss are largely restricted to repeat-rich regions, particularly ERVs, and shows little association with activation of the male transcriptional program. These results indicate that changes in chromatin accessibility and H3K27ac, rather than H3K9me3 loss, represent the principal chromatin alterations associated with granulosa-to-Sertoli-like transdifferentiation.

### TRIM28 is frequently associated with the ovarian transcription factor network

The enrichment of ovarian transcription factor motifs within TRIM28 binding sites and in regions that lose accessibility upon TRIM28 deletion prompted us to examine more closely the regulatory network centered on FOXL2, NR5A2 (a.k.a. LHR-1), ESR2 and RUNX1. These factors contribute to granulosa cell identity, maturation and function (2, 23, 26, 36–41), and co-localization of FOXL2 with TRIM28, NR5A2 and ESR2 on chromatin have been previously shown using ChIP-seq (24) and ChIP-SICAP (42). To investigate how these transcription factors are associated with TRIM28 in the adult ovary, we reanalyzed publicly available ChIP-seq datasets for these transcription factors and assessed the overlap of their binding sites with TRIM28 occupancy from control ovaries.

We found extensive co-occupancy between TRIM28 and all four transcription factors (Fig. 4a). Approximately half of all FOXL2-bound regions (49.4%) and more than half of NR5A2-bound regions (55.5%) were also occupied by TRIM28. Co-occupancy was even more pronounced for RUNX1, with 71.2% of RUNX1 binding sites overlapping TRIM28 peaks. Although lower, ESR2 binding sites also showed substantial overlap with TRIM28 (19.5%). Genome-wide permutation tests confirmed that all overlaps were significantly greater than expected by chance given their repartition on the genome, with enrichments ranging from 10.7-fold for ESR2 to 32.6-fold for RUNX1 (Fig. 4a). These results indicate that TRIM28 is broadly integrated within the ovarian transcription factor network and occupies a substantial fraction of regulatory regions bound by key granulosa cell transcription factors.

**Figure 4:**
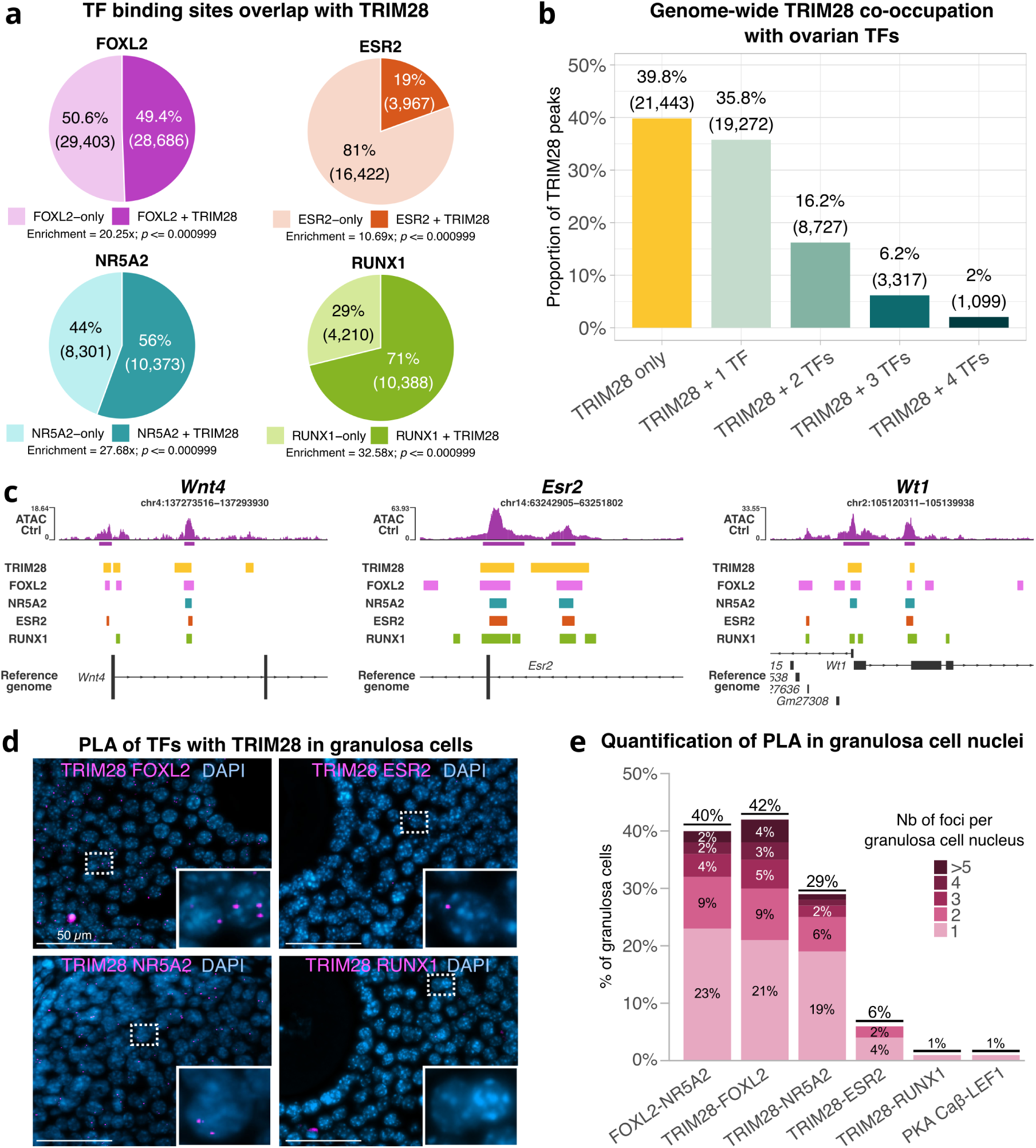
TRIM28 is extensively integrated into the ovarian transcription factor network. (**a**) Pie charts showing the proportion of FOXL2, NR5A2, ESR2, and RUNX1 binding sites co-occupied by TRIM28 in adult control ovaries. Fisher’s exact test p-values (TF and TRIM28 colocalization enrichment compared to random genomic distribution) are indicated below each chart. (**b**) Distribution of TRIM28-bound regions according to the number of co-occupying ovarian TFs. (**c**) Genome browser tracks showing examples of TRIM28+TF co-occupancy at key ovarian loci. (**d**) Microscopy images of proximity ligation assays (PLA) of the four ovarian TFs with TRIM28 in granulosa cell nuclei from 8-week-old control ovaries. (**e**) Quantification of different PLA in granulosa cell nuclei from 8-week-old control ovaries. The PKA Capβ-LEF1 PLA is a negative control.

To further characterize the relationship between TRIM28 and the ovarian transcription factor network, we classified TRIM28-bound regions according to the number and identity of co-occupying ovarian transcription factors (Fig. 4b, Supplementary Fig. S6, Supplementary Table S7). Overall, 60.2% of TRIM28 peaks overlapped at least one of the four ovarian transcription factors analyzed (Fig. 4b). Among these regions, 35.8% were co-occupied by one transcription factor, whereas 16.2%, 6.2% and 2% were co-occupied by two, three or four of them, respectively. These results indicate that a substantial proportion of TRIM28 binding sites correspond to regulatory regions occupied by multiple ovarian transcription factors. Analysis of the most frequent transcription factor combinations revealed a strong enrichment for FOXL2- and NR5A2-containing modules (Supplementary Fig. S6). In total, 1,099 TRIM28-bound regions were occupied by all four ovarian transcription factors (Fig. 4b, Supplementary Table S6), including regulatory regions associated with the *Wnt4*, *Wt1* and *Esr2* loci (Fig. 4c).

To determine whether the genomic co-occupancy identified by chromatin profiling reflects close spatial proximity in granulosa cells, we performed proximity ligation assays (PLA) between TRIM28 and each of the four ovarian transcription factors (Fig. 4d-e). Expression of all proteins was first confirmed in 8-week-old ovaries using the antibodies employed for PLA (Supplementary Fig. S7). PLA confirmed the previously reported proximity between FOXL2 and NR5A2, as well as between FOXL2 and TRIM28 (42), with signals detected in 40% and 42% of granulosa cell nuclei, respectively (Fig. 4e). A robust PLA signal was also observed for the TRIM28-NR5A2 pair, with 29% of granulosa cells displaying at least one PLA focus, consistent with the extensive overlap of their genomic binding sites (Fig. 4a). By contrast, only a small fraction of granulosa cells showed a PLA signal between TRIM28 and ESR2, consistent with their more limited genomic co-occupancy. Surprisingly, despite the substantial overlap between TRIM28 and RUNX1 binding sites, no significant PLA signal was detected between TRIM28 and RUNX1. These results suggest that although both proteins occupy the same regulatory regions, they are unlikely to be present within the <40 nm distance required for PLA detection.

Together, these results indicate that TRIM28 is closely associated with the ovarian transcription factor network and preferentially occupies regulatory regions bound by multiple ovarian transcription factors.

### TRIM28-associated ovarian regulatory regions preferentially lose chromatin accessibility following *Trim28* deletion

Because TRIM28 extensively co-occupies ovarian regulatory regions FOXL2, NR5A2, ESR2 and RUNX1, we next asked whether these regions were preferentially affected by *Trim28* deletion. To address this question, we examined the distribution of TRIM28 and ovarian transcription factor binding sites across regions that either lost (Fig. 3a, cluster a) or gained (cluster b) chromatin accessibility in *Trim28* cKO ovaries.

Regions losing accessibility were strongly enriched for TRIM28 occupancy and ovarian transcription factor binding, whereas regions gaining accessibility showed much lower overlap with these factors (Fig. 5a). This suggests that these transcription factors primarily contribute to the maintenance of ovarian regulatory elements rather than to the repression of regions that become activated following *Trim28* deletion. Among TRIM28-bound regions, the proportion of regions losing accessibility increased progressively with the number of co-occupying ovarian transcription factors, from 3.1% at sites bound by TRIM28 alone to 5.1%, 7.9%, 12.9% and 20% at sites co-occupied by one, two, three or four ovarian transcription factors, respectively (Fig. 5a). Accordingly, TRIM28-bound regions occupied by at least two ovarian transcription factors were significantly enriched among regions losing accessibility and depleted among regions gaining accessibility (Fisher’s exact test, odd ratio = 2.02, *p*-value < 2.2×10^-16^). Representative examples are shown in Fig. 5b. Regulatory regions associated with *Foxl2*, *Esr2*, *Emx2* and *Bmp2* were occupied by TRIM28 together with all four ovarian transcription factors and showed reduced chromatin accessibility following *Trim28* deletion. Together, these results indicate that regulatory regions co-occupied by TRIM28 and multiple ovarian transcription factors are particularly dependent on TRIM28 for the maintenance of chromatin accessibility.

**Figure 5:**
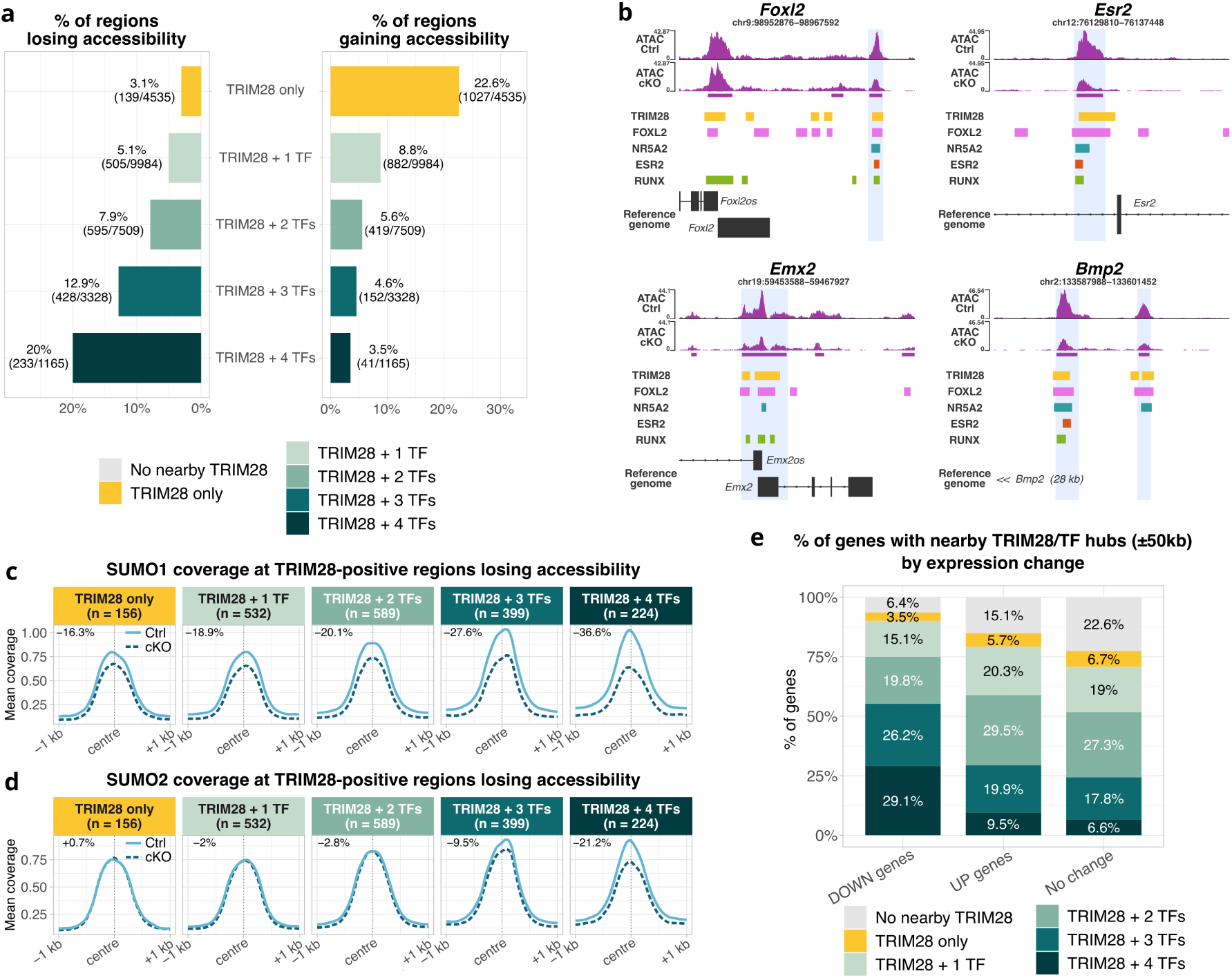
Loss of TRIM28 destabilizes multi-TF regulatory hubs and downregulates ovarian genes. (**a**) Distribution of TRIM28-bound chromatin regions that lose (left) or gain (right) accessibility in *Trim28* cKO ovaries, classified by the number of co-occupying ovarian transcription factors (FOXL2, NR5A2, ESR2, RUNX). (**b**) Genome browser tracks showing examples of ovarian regulatory hubs where TRIM28 and all four ovarian TFs co-localize and exhibit decreased accessibility in *Trim28* cKO. (**c**, **d**) SUMO1 (**c**) and SUMO2 (**d**) ChIP-seq coverage at TRIM28-bound regions losing accessibility in *Trim28* cKO ovaries, stratified by ovarian TF co-occupancy. Solid light-blue lines: control (Ctrl); dashed dark-blue lines: *Trim28* cKO. Percentage of signal change (Ctrl vs. cKO) is indicated in the upper-left corner of each panel. (**E**) Association between gene expression changes (scRNA-seq from Rossitto et al., 2022) and TRIM28+TF co-occupancy within ±50 kb of gene bodies. Genes are classified as downregulated (DOWN), upregulated (UP), or unchanged upon *Trim28* cKO.

As TRIM28 extensively co-occupies regulatory regions with ovarian transcription factors, we next asked whether the expression of these factors was altered following *Trim28* deletion. Analysis of our previously published single-cell RNA-seq dataset (7) revealed that *Foxl2* and *Nr5a2* expression was already reduced in granulosa cells from *Trim28* cKO ovaries and was retained in only a small fraction of Sertoli-like cells (Supplementary Fig. S8). By contrast, *Esr2* expression was maintained in granulosa cells but decreased in Sertoli-like cells, whereas *Runx1* expression remained largely unchanged. These observations suggest a progressive destabilization of the ovarian transcriptional program, whereby the initial loss of TRIM28 weakens expression of key ovarian transcription factors, which may further destabilize the regulatory circuitry required to maintain granulosa cell identity.

Because SUMOylation is a key molecular function of TRIM28, we reasoned that impaired SUMOylation might contribute to the progressive destabilization of the ovarian transcriptional program. We therefore asked whether SUMOylation at these regulatory regions was altered following *Trim28* deletion. To this aim, we reanalyzed the SUMO1 and SUMO2 ChIP-seq datasets previously generated in control and *Trim28* cKO ovaries (7). SUMO1 and SUMO2 deposition were reduced at TRIM28-bound DARs that lost chromatin accessibility following *Trim28* deletion (Fig. 5c-d). Moreover, the extent of SUMO loss increased progressively with the number of ovarian transcription factors co-occupying TRIM28-bound regulatory regions (Spearman’s correlation = 0.99 for both SUMO1 and SUMO2), with the greatest reduction observed at sites co-occupied by TRIM28 and all four ovarian transcription factors (36.6% and 21.2% reduction of SUMO1 and SUMO2, respectively, Fig. 5c-d). Together, these results indicate that the TRIM28-bound regions that are most dependent on TRIM28 for maintaining chromatin accessibility also undergo the greatest loss of SUMOylation following *Trim28* deletion.

We next asked whether genes whose expression changed following *Trim28* deletion (8-week-old ovaries control and cKO single-cell RNA-seq data) (7) were preferentially associated with TRIM28-bound regulatory regions occupied by multiple ovarian transcription factors. To address this question, each expressed gene was assigned to the TRIM28-associated region located within ±50 kb that co-localize with the greatest number of co-occupying ovarian transcription factors (from TRIM28 alone to TRIM28 with all four transcription factors). We then compared the distribution of these TF hub classes among downregulated, upregulated and non-differentially expressed genes (Fig. 5e).

Downregulated genes were strongly enriched for TRIM28 regions co-occupied with several TF. Nearly one third of downregulated genes (29.1%) were associated with a nearby TRIM28 region occupied by all four ovarian transcription factors, compared with only 6.6% of non-differentially expressed genes (Fig. 5e). Similarly, 55.3% of downregulated genes were associated with TRIM28 regions occupied by at least three ovarian transcription factors, compared with 24.4% of non-differentially expressed genes. Fisher’s exact tests confirmed a strong enrichment of downregulated genes among loci associated with TRIM28 regions co-occupied by at least three transcription factors (odds ratio = 3.8, p < 2.2×10^-^¹⁶) and, more specifically, among loci associated with TRIM28 regions occupied by all four transcription factors (odds ratio = 5.74, p < 2.2×10^-^¹⁶) (Fig. 5e). In contrast, upregulated genes showed only modest enrichment for these highly co-occupied regulatory regions (odds ratio = 1.28 and 1.46 for ≥3 TFs and 4 TFs, respectively). Together, these results indicate that genes downregulated following *Trim28* deletion are preferentially associated with TRIM28-bound regulatory regions co-occupied by multiple ovarian transcription factors.

Overall, these results show that *Trim28* deletion preferentially affects regulatory regions co-occupied by multiple ovarian transcription factors, resulting in reduced chromatin accessibility, decreased SUMOylation and downregulation of nearby genes.

## DISCUSSION

Maintenance of ovarian identity requires continuous repression of the testicular differentiation program after birth. We have previously shown that conditional deletion of *Trim28* or disruption of its PHD-dependent SUMO E3 ligase activity induces progressive granulosa-to-Sertoli transdifferentiation (7). Because TRIM28 functions both as a regulator of H3K9me3-dependent heterochromatin and as a transcriptional activator, the relative contribution of these activities to ovarian maintenance remained unclear. Here, we show that these functions contribute differently to ovarian identity. While the canonical TRIM28-SETDB1 pathway primarily maintains repeat silencing, our data identify TRIM28 activity at cis-regulatory elements as the principal mechanism preserving granulosa cell identity.

In most cellular contexts, TRIM28 is recruited by KRAB zinc-finger proteins to repetitive sequences, where it promotes SETDB1- and HP1-dependent H3K9me3 deposition and heterochromatin formation (9, 10, 43). Accordingly, 30-45% of TRIM28 binding sites overlap H3K9me3 domains in several mammalian cell types (35, 44–46). In contrast, we found that ovarian TRIM28 predominantly occupies promoters and distal regulatory elements enriched for ovarian transcription factor motifs, with only a small proportion of binding sites associated with H3K9me3 (13.6%). These observations indicate that, in ovarian somatic cells, TRIM28 acts predominantly at cis-regulatory elements rather than through its canonical heterochromatin-associated function. Consistent with this, *Trim28* deletion produced only limited remodeling of the H3K9me3 landscape. Most H3K9me3 domains remained unchanged, suggesting that their maintenance relies on redundant mechanisms involving additional H3K9 methyltransferases, HP1 proteins and DNA methylation (46, 47).

H3K9me3 loss showed little association with testis-determining genes, Sertoli-specific regulatory elements or testicular transcription factor motifs. In particular, we found no evidence of H3K9me3 loss at the regulatory regions of the key testis-determining genes *Sox9* and *Dmrt1*, despite their transcriptional activation in *Trim28* cKO ovaries. Instead, it mainly affected repeat-rich regions, particularly ERVs, consistent with the established role of the TRIM28-SETDB1 pathway in transposable element repression (9, 15). Although endogenous retroviruses can function as alternative promoters or enhancers (34), we found no evidence that ERVs losing H3K9me3 contribute to activation of the male transcriptional program (no known upregulated testis genes nearby). These observations indicate that H3K9me3 remodeling alone is unlikely to drive granulosa-to-Sertoli transdifferentiation. Finally, H3K9me3 depletion at loci lacking detectable TRIM28 binding in adult ovaries may reflect earlier developmental functions of TRIM28 that are no longer apparent at this stage. Temporal chromatin profiling will be required to test this possibility.

By contrast, changes in chromatin accessibility and H3K27ac closely paralleled the transcriptional changes observed in *Trim28* cKO ovaries. Regions losing accessibility were enriched for ovarian transcription factor motifs, whereas newly accessible regions were enriched for SOX motifs but not DMRT1 motifs. Given that ectopic DMRT1 expression is sufficient to induce granulosa-to-Sertoli transdifferentiation (6), the absence of DMRT1 motif enrichment suggests that the establishment of the male regulatory program in *Trim28* cKO ovaries is not primarily driven by DMRT1.

Remarkably, when looking at regions originally bound by TRIM28 in controls, the greatest reductions in chromatin accessibility occurred at regulatory regions co-occupied by TRIM28 and multiple ovarian transcription factors. These regions were also associated with decreased expression of nearby genes. Together, these findings indicate that TRIM28 preferentially maintains the activity of regulatory regions co-occupied by multiple ovarian transcription factors, supporting the expression of the ovarian transcriptional program.

Our data also provides insight into the relationship between TRIM28, ovarian transcription factors and SUMOylation. Previous studies, including our own, showed that TRIM28 co-localizes with FOXL2 on ovarian chromatin and that disruption of its PHD-dependent SUMO E3 ligase activity is sufficient to induce ovarian masculinization (7, 42). Here, we extend these observations by showing that TRIM28 also extensively co-occupies regulatory regions with FOXL2, NR5A2, ESR2 and RUNX1, and that these regions are particularly affected by *Trim28* deletion. In addition, SUMO1 and SUMO2 deposition was progressively reduced at TRIM28-bound regulatory regions as the number of co-occupying ovarian transcription factors increased. Together, these findings suggest that TRIM28-dependent SUMOylation preferentially occurs at regulatory regions co-occupied by multiple ovarian transcription factors, where it may contribute to maintaining their activity.

However, the identity of the SUMOylated proteins at these regions remains unknown. Although histone SUMOylation has been described, it is generally a low-abundance and highly dynamic modification associated primarily with transcriptional repression (48), making it unlikely to account for the coordinated loss of chromatin accessibility at these active regulatory elements. Instead, the SUMO signal is likely to reflect SUMOylation of chromatin-associated transcription factors. This is supported by the preferential loss of SUMOylation at regions co-occupied by FOXL2, NR5A2, ESR2 and RUNX1, together with our previous finding that TRIM28 promotes SUMOylation of FOXL2 and RUNX1 (7) and reports showing that SUMOylation stabilizes FOXL2 (49), ESR2 (50) and RUNX1 (51). Together, these findings suggest that TRIM28-dependent SUMOylation contributes to the stability and/or chromatin occupancy of ovarian transcription factors required to maintain the activity of ovarian regulatory elements.

Based on these findings, we propose that loss of TRIM28 progressively destabilizes the ovarian transcriptional program. The *Foxl2*, *Nr5a2* and *Esr2* loci contain candidate regulatory elements co-occupied by TRIM28 and multiple ovarian transcription factors, suggesting that TRIM28 contributes to maintaining their expression. Reduced stability of ovarian transcription factors resulting from the loss of TRIM28 E3 SUMO ligase activity at these regulatory elements in mutant would in turn decrease the expression of key ovarian transcription factors, further compromising the activity of their shared regulatory regions and progressively promoting granulosa-to-Sertoli transdifferentiation.

Although our genomic and PLA analyses indicate that TRIM28 is closely associated with multiple ovarian transcription factors, they do not establish whether FOXL2, ESR2, NR5A2 and RUNX1 are simultaneously associated with TRIM28 on chromatin. It is therefore possible that distinct transcription factors interact with TRIM28 sequentially during folliculogenesis. Determining whether TRIM28 assembles a single multiprotein complex or instead forms distinct dynamic complexes will require future biochemical studies.

Unlike previously described TRIM28-deficient systems, in which transposable element activation (35) and heterochromatin disruption (52) dominate the phenotype, our data indicate that disruption of lineage-specific cis-regulatory elements represents the major consequence of TRIM28 loss in ovarian somatic cells. Together, these findings suggest that the relative contribution of TRIM28 to heterochromatin maintenance and transcriptional regulation is highly cellular context dependent and reflects the genomic distribution of its lineage-specific binding partners.

An unresolved question concerns the mechanism underlying the increased chromatin accessibility at testis-specific regulatory regions following *Trim28* deletion. Although many of these regions are bound by TRIM28 in control ovaries, only a minority overlaps ovarian transcription factor binding sites. Our data indicate that this chromatin opening is unlikely to result from H3K9me3 loss, which primarily affects ERVs. Moreover, a direct effect through H3K27me3 appears unlikely, as TRIM28 is not known to recruit PRC2 despite reported interactions with EZH2 that primarily promote transcriptional activation rather than PRC2-mediated gene repression (53). By contrast, TRIM28 has been shown to recruit PRC1 to differentiation-associated genes in embryonic stem cells (54) and to stabilize the PRC1 component CBX8 (55). We therefore speculate that *Trim28* deletion may impair PRC1-dependent H2AK119ub deposition, thereby facilitating chromatin opening at testis regulatory elements. Future studies will be required to determine whether TRIM28 contributes to PRC1 occupancy and H2AK119ub deposition at these loci.

In conclusion, our findings support a model in which TRIM28 maintains ovarian identity through two complementary genomic functions. While it contributes to heterochromatin organization and repression of repetitive elements, its predominant role in the adult ovary is to preserve the integrity of ovarian transcription factor hubs via its E3 SUMOligase activity. Destabilization of these hubs progressively weakens the granulosa transcriptional program, leading to the collapse of the ovarian regulatory network and ultimately permitting granulosa-to-Sertoli transdifferentiation. Our findings therefore identify TRIM28-dependent SUMOylation as a central organizer of the lineage-specific TF hubs that ensure long-term maintenance of ovarian identity.

## Supporting information

SupFigures

## ACKNOWLEDGEMENTS

We are grateful to the genotoul bioinformatics platform Toulouse Occitanie (Bioinfo Genotoul, https://doi.org/10.15454/1.5572369328961167E12) for providing the computing resources necessary for the current study. We thank members of our lab for advice, support, and helpful comments. We are grateful to the staff (particularly Clarisse Froget) of the IGH animal care facility. We also thank the staff of IGH Imaging facility (MRI Montpellier) for their help with microscopy analyses.

## AUTHOR CONTRIBUTION

Conceptualization: L.S., I.S. and F.P. Investigation: L.S., F.Ch., G.G., S.D., F.Ca., A.-M.L.-M., I.S., and F.P. Methodology: L.S., F.Ch., G.G., S.D., F.Ca., A.M.L.M., I.S. and F.P. Data curation: I.S. and F.P. Formal analysis: I.S. and F.P. Visualization: M.R. and I.S. Software: I.S. Resources: F.Ch., G.G., S.D., F.Ca., A.-M.L.-M., I.S., and F.P. Writing-review and editing: L.S., F.Ch., G.G., S.D., F.Ca., A.-M.L.-M., I.S., and F.P. Project administration: I.S. and F.P. Supervision: A.-M.L.-M.,I.S., and F.P. Funding acquisition: A.-M.L.-M., I.S. and F.P. All authors have read and accepted the data being presented in the manuscript.

## COMPETING DECLARATION

The authors declare that they have no competing interests.

## DATA AVAILABILITY

CUT&RUN data for H3K9me3 and H3K27ac, as well as ATAC-seq data produced in the current study have been deposited in the Gene Expression Omnibus under accession numbers GSE339481 and GSE339478, respectively. The data is only accessible to reviewers at the moment.

UCSC genome browser session: https://genome.ucsc.edu/s/IStevant/TRIM28_mouse_ovary

The code produced to analyze the data is available on Zenodo: https://zenodo.org/records/21493393

