## Supplementary material for "TRIM28 PRESERVES OVARIAN IDENTITY BY STABILIZING LINEAGE-SPECIFIC TRANSCRIPTION FACTOR HUBS": SupFigures

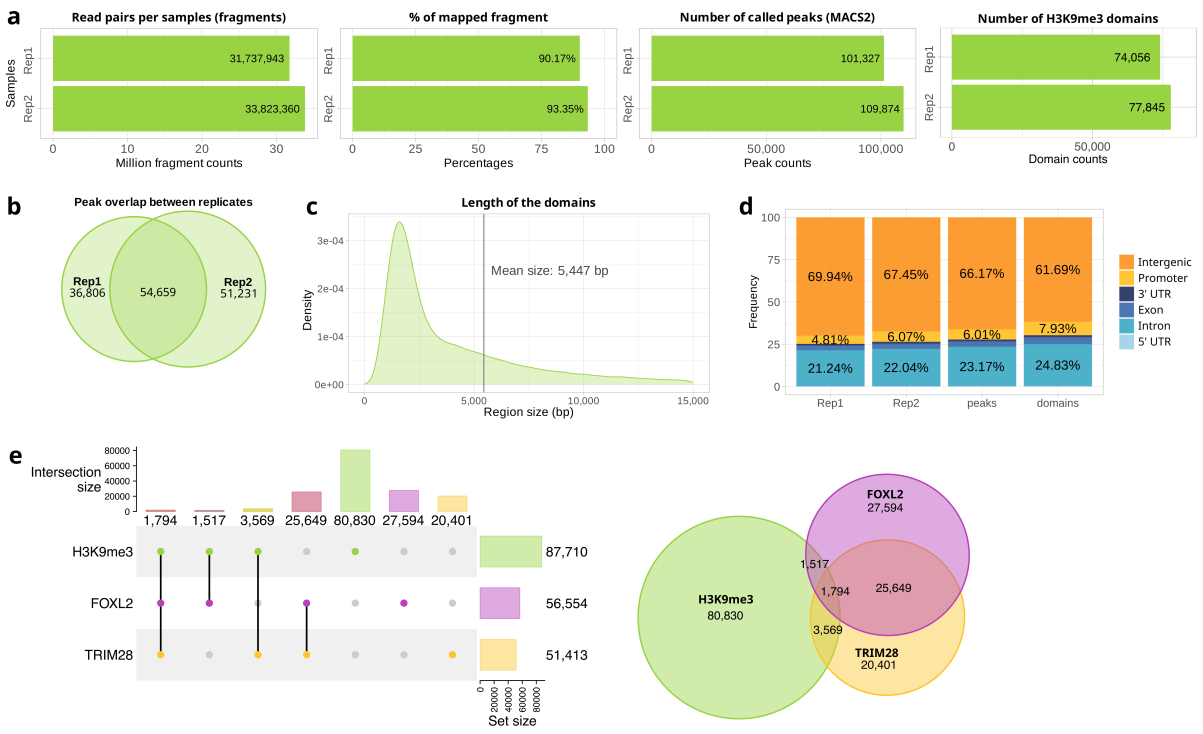


**Supplementary Figure S1**: Quality controls and characteristics of H3K9me3 CUT&RUN in control ovaries. (**a**) Sequencing statistics for the two biological replicates, showing the number of read pairs, mapping rate, number of MACS2 peaks and number of merged H3K9me3 domains. (**b**) Overlap between H3K9me3 peaks identified in the two biological replicates. (**c**) Distribution of H3K9me3 domain lengths after merging peaks separated by less than 2 kb. The mean domain size is indicated. (**d**) Genomic annotation of H3K9me3 peaks in each replicate and in the consensus datasets. (**e**) UpSet plot (left) and Venn diagram (right) showing the overlap between H3K9me3 domains, TRIM28 and FOXL2 ChIP-seq peaks in control ovaries.

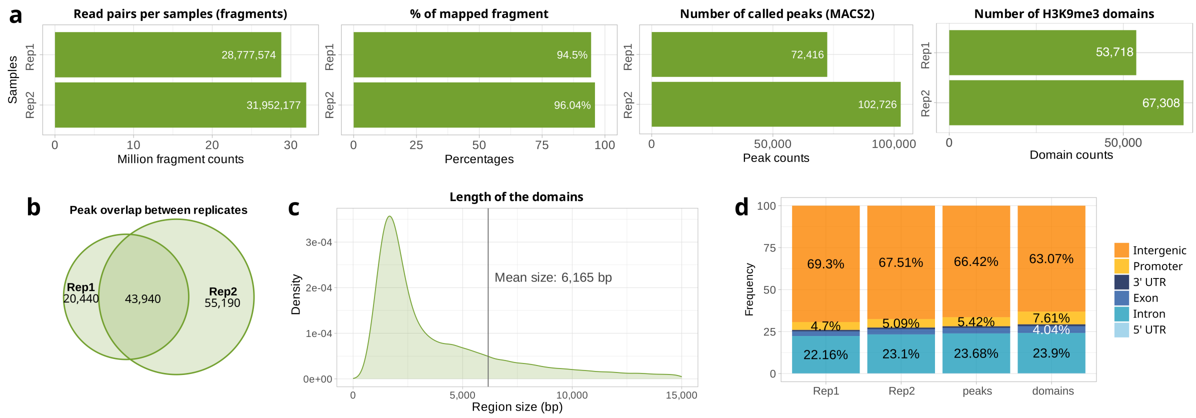


**Supplementary Figure S2:** Quality controls and characteristics of H3K9me3 CUT&RUN in *Trim28* cKO ovaries. (**a**) Sequencing statistics for the two biological replicates, showing the number of read pairs, mapping rate, number of MACS2 peaks and number of merged H3K9me3 domains. (**b**) Overlap between H3K9me3 peaks identified in the two biological replicates. (**c**) Distribution of H3K9me3 domain lengths after merging peaks separated by less than 2 kb. (**d**) Genomic annotation of H3K9me3 peaks in each replicate and in the consensus datasets.

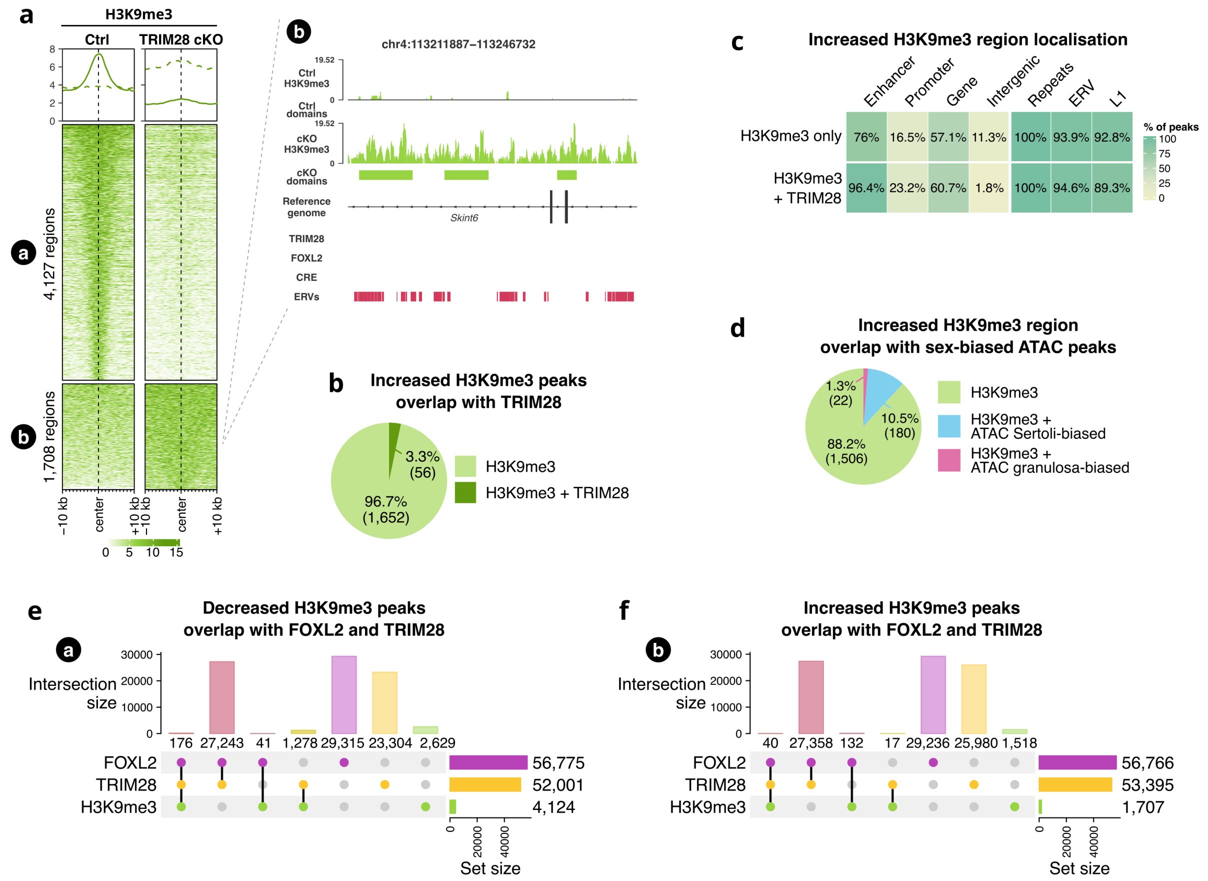


**Supplementary Figure S3**: Characterisation of genomic regions gaining H3K9me3 following *Trim28* deletion. (**a**) Heatmap of H3K9me3 signal at regions showing increased H3K9me3 in *Trim28* cKO ovaries compared with controls. Right panel: representative genome browser view showing increased H3K9me3 signal together with TRIM28 and FOXL2 ChIP-seq tracks, cis-regulatory elements (CREs) and ERVs. (**b**) Proportion of H3K9me3-gained regions overlapping TRIM28 binding sites (±1 kb). (**c**) Genomic annotation of H3K9me3-gained regions with or without nearby TRIM28 binding. (**d**) Overlap between H3K9me3-gained regions and previously described granulosa- and Sertoli-biased open chromatin regions. (**e,f**) UpSet plots showing the overlap between FOXL2, TRIM28 and H3K9me3 regions that lose (**e**) or gain (**f**) H3K9me3 following *Trim28* deletion.

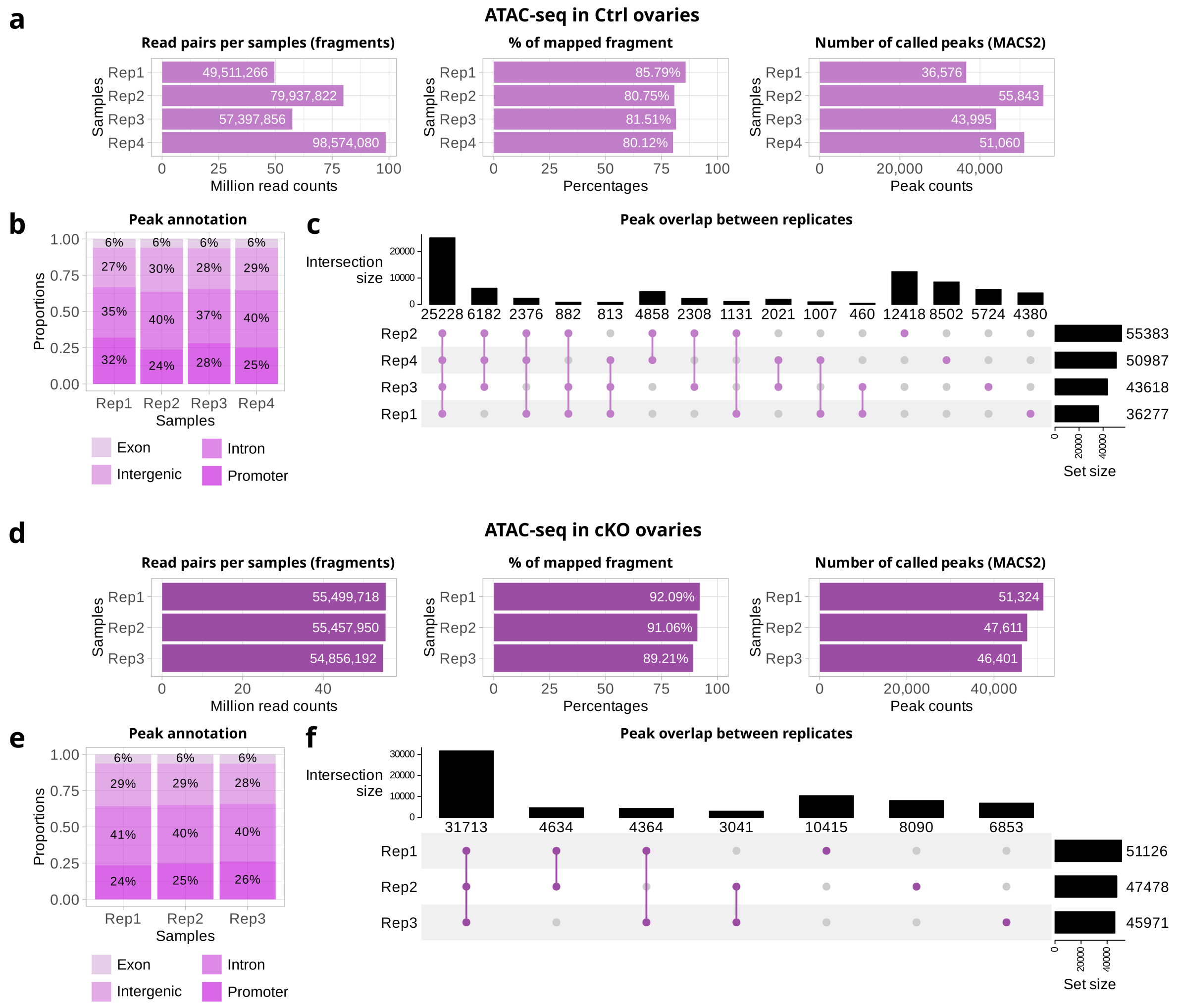


**Supplementary Figure S4:** Quality controls for ATAC-seq datasets. (**a-c**) Quality control metrics for ATAC-seq performed on control ovaries, including sequencing statistics, genomic annotation of peaks and overlap between biological replicates. (**d-f**) Corresponding quality control analyses for *Trim28* cKO ovaries.

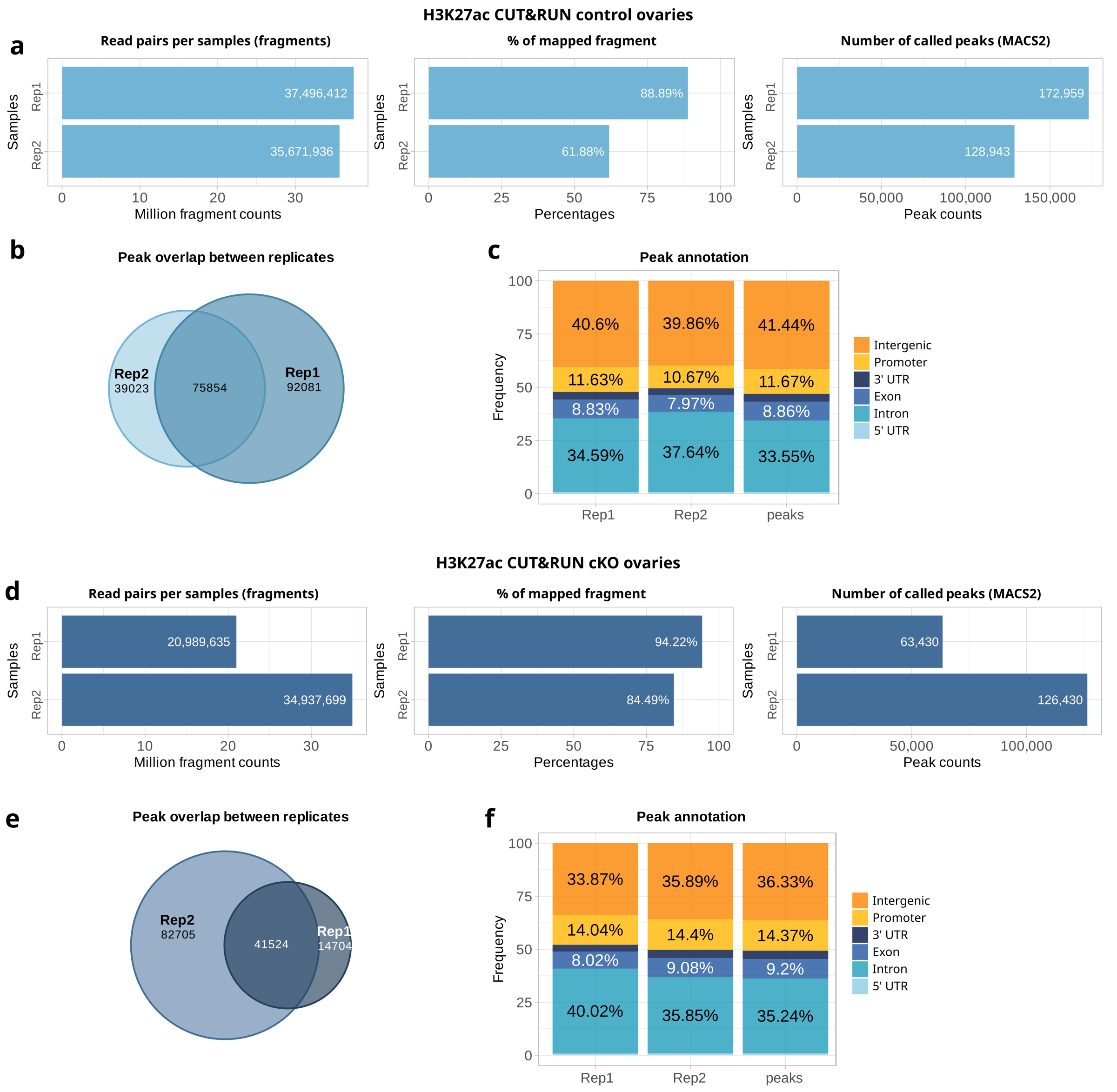


**Supplementary Figure S5:** Quality controls for H3K27ac CUT&RUN datasets. (**a-c**) Sequencing statistics, overlap between biological replicates and genomic annotation of H3K27ac peaks in control ovaries. (**d-f**) Corresponding quality control analyses for H3K27ac CUT&RUN performed in *Trim28* cKO ovaries.

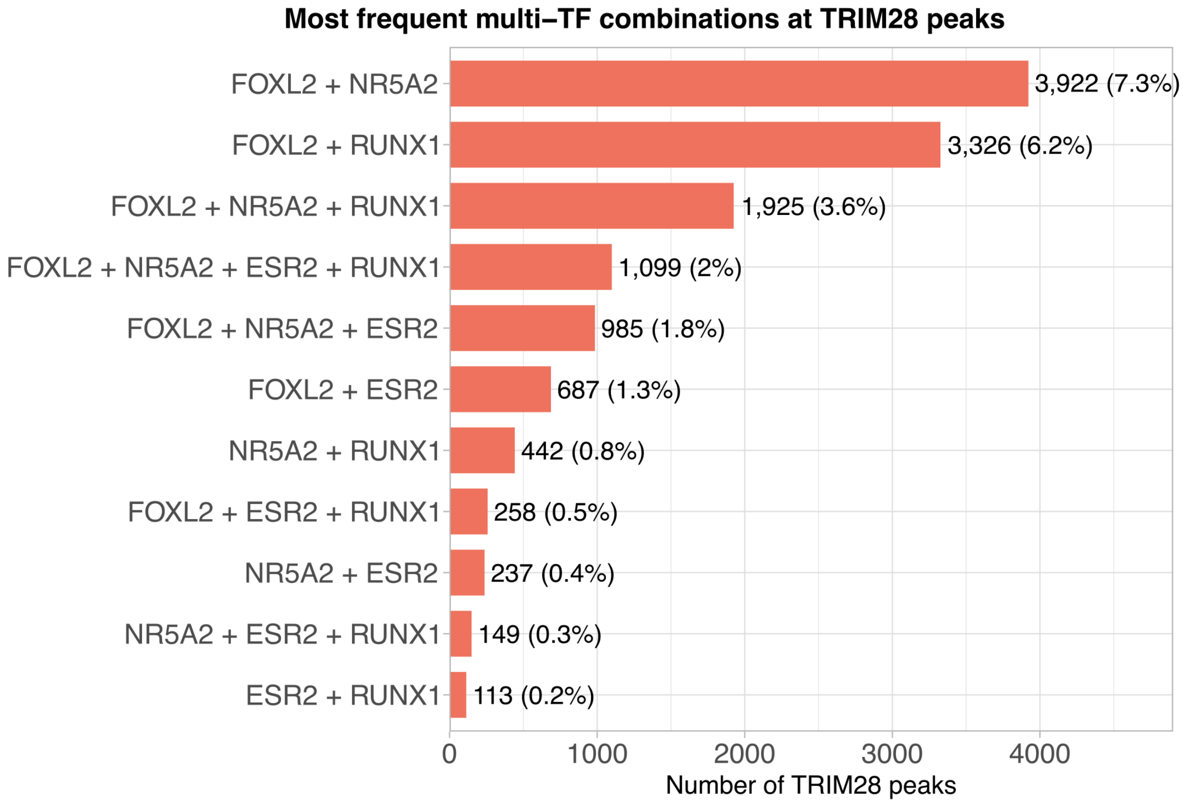


**Supplementary Figure S6:** Most frequent combinations of ovarian transcription factors at TRIM28-bound regulatory regions. Frequency of the most abundant combinations of FOXL2, NR5A2, ESR2 and RUNX1 binding at TRIM28-associated regulatory regions. Bars indicate the number and percentage of TRIM28 peaks co-occupied by each transcription factor combination.

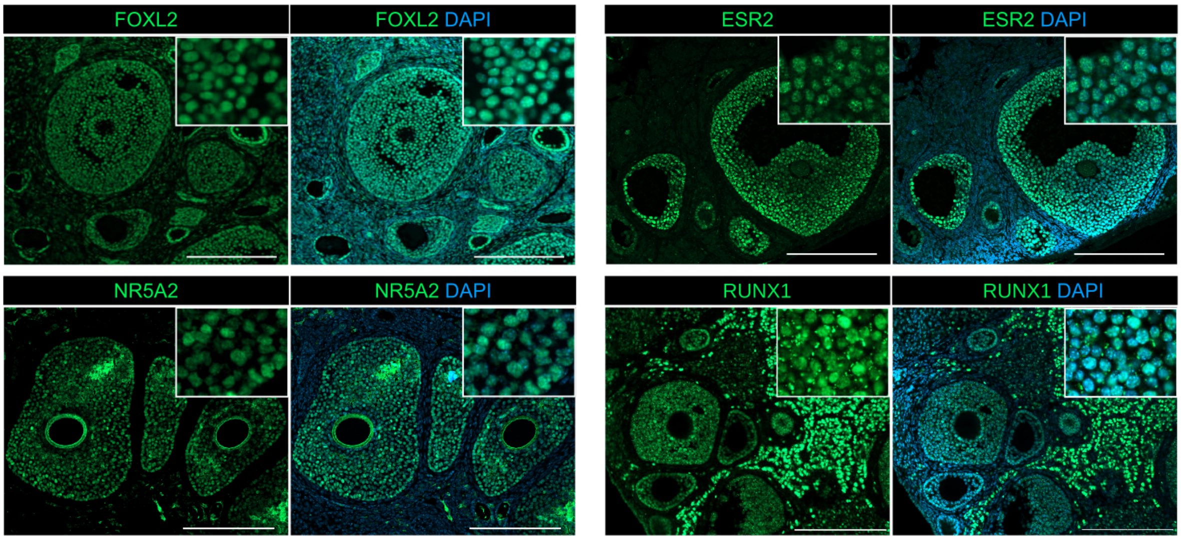


**Supplementary Figure S7:** Expression of ovarian transcription factors in adult control ovaries. Representative immunofluorescence images showing expression of FOXL2, ESR2, NR5A2 and RUNX1 in granulosa cells of adult control ovaries. Nuclei are counterstained with DAPI. Insets show higher-magnification views of granulosa cell nuclei.

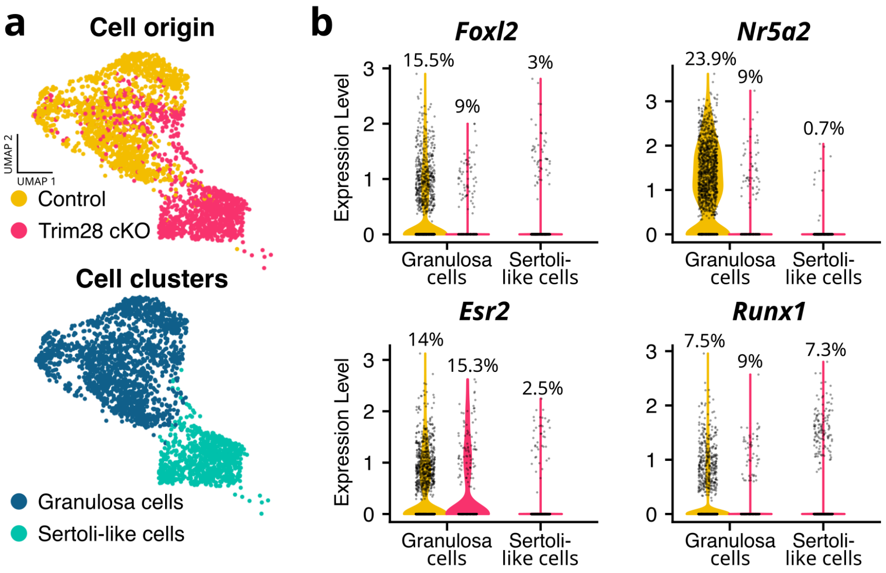


**Supplementary Figure S8:** Expression of ovarian transcription factors during granulosa-to-Sertoli-like transdifferentiation in *Trim28* cKO ovaries. (**a**) UMAP representation of the previously published single-cell RNA-seq dataset from control and Trim28 cKO adult ovaries, coloured by genotype (top) or cell identity (bottom). Granulosa cells comprise both control and remaining *Trim28* cKO granulosa cells, whereas Sertoli-like cells are exclusively derived from *Trim28* cKO ovaries. (**b**) Violin plots showing expression of *Foxl2*, *Nr5a2*, *Esr2* and *Runx1* in granulosa and Sertoli-like cells. Violin colours indicate cell origin (control, yellow; *Trim28* cKO, pink). Individual cells are shown as dots.

**Supplementary Table S1:** List of the antibodies used in the study.

**Supplementary Table S2:** H3K9me3 landscape in control ovaries. This table contains the genomic coordinates, annotation and associated analyses of H3K9me3-enriched regions identified by CUT&RUN in control ovarian somatic cells.

Worksheets:

- Ctrl H3K9me3 domains: genomic coordinates and annotation of merged H3K9me3 domains.

- Ctrl H3K9me3 peaks: genomic coordinates and annotation of individual H3K9me3 peaks.

- GO terms genes in H3K9me3: Gene Ontology enrichment analysis of genes associated with H3K9me3 domains.

- Raw read counts: raw fragment counts for all H3K9me3 regions used for differential analysis.

**Supplementary Table S3:** TRIM28 and FOXL2 genomic binding sites. This table contains the genomic coordinates and annotations of TRIM28 and FOXL2 ChIP-seq peaks together with overlap analyses used throughout the study.

Worksheets:

- TRIM28 peaks: genomic coordinates and annotation of TRIM28 ChIP-seq peaks.

- FOXL2 peaks: genomic coordinates and annotation of FOXL2 ChIP-seq peaks.

- TRIM28 + FOXL2 overlap: genomic regions co-occupied by TRIM28 and FOXL2.

**Supplementary Table S4:** Transcription factor motif enrichment at TRIM28-associated chromatin. This table contains transcription factor binding site (TFBS) motif enrichment analyses performed on TRIM28-associated chromatin compartments.

Worksheets:

- TRIM28-only TFBS: motif enrichment in TRIM28-only peaks.

- TRIM28 + H3K9me3 TFBS: motif enrichment in regions co-occupied by TRIM28 and H3K9me3.

- H3K9me3-only TFBS: motif enrichment in H3K9me3-only regions.

**Supplementary Table S5:** Differential H3K9me3 regions following Trim28 deletion. This table contains genomic regions showing significant changes in H3K9me3 following Trim28 deletion together with their genomic annotation, associated genes and downstream analyses.

Worksheets:

- H3K9me3 DER summary: genomic coordinates, annotation and overlap analyses of differentially enriched H3K9me3 regions.

- H3K9me3 DER statistics: DESeq2 differential enrichment statistics.

- GO terms genes in DER H3K9me3: Gene Ontology enrichment analysis of genes associated with differentially enriched H3K9me3 regions.

- H3K9me3 DER TFBS: transcription factor motif enrichment analysis of regions showing H3K9me3 loss or gain.

- Raw read counts: raw fragment counts for all regions included in the differential analysis.

**Supplementary Table S6:** Chromatin accessibility changes following Trim28 deletion. This table contains ATAC-seq peaks, differentially accessible regions (DARs), motif enrichment analyses and transposable element annotations.

Worksheets:

- Ctrl ATAC peaks: accessible chromatin regions identified in control ovaries.

- cKO ATAC peaks: accessible chromatin regions identified in Trim28 cKO ovaries.

- DAR ATAC summary: genomic coordinates, cluster assignment, genomic annotation and overlap analyses of differentially accessible regions.

- DAR ATAC statistics: DESeq2 differential accessibility statistics.

- DAR ATAC TFBS: transcription factor motif enrichment analyses for regions losing (cluster a) or gaining (cluster b) chromatin accessibility.

- DE TEs in DARs: differentially expressed transposable elements overlapping differentially accessible regions.

- Raw read counts: raw fragment counts used for differential accessibility analysis.

**Supplementary Table S7:** Co-occupancy of ovarian transcription factors at TRIM28-bound regulatory regions. This table contains the genomic coordinates of TRIM28-bound regulatory regions together with the presence or absence of FOXL2, NR5A2, ESR2 and RUNX1 binding, the total number of co-occupying transcription factors, and the corresponding transcription factor combination.
